# Identification of FacZ as a division site placement factor in *Staphylococcus aureus*

**DOI:** 10.1101/2023.04.24.538170

**Authors:** Thomas M. Bartlett, Tyler A. Sisley, Aaron Mychack, Suzanne Walker, Richard W. Baker, David Z. Rudner, Thomas G. Bernhardt

## Abstract

*Staphylococcus aureus* is a gram-positive pathogen responsible for life-threatening infections that are difficult to treat due to antibiotic resistance. The identification of new vulnerabilities in essential processes like cell envelope biogenesis represents a promising avenue towards the development of anti-staphylococcal therapies that overcome resistance. To this end, we performed cell sorting-based enrichments for *S. aureus* mutants with defects in envelope integrity and cell division. We identified many known envelope biogenesis factors as well as a large collection of new factors with roles in this process. Mutants inactivated for one of the hits, the uncharacterized SAOUHSC_01855 protein, displayed aberrant membrane invaginations and multiple FtsZ cytokinetic ring structures. This factor is broadly distributed among Firmicutes, and its inactivation in *B. subtilis* similarly caused division and membrane defects. We therefore renamed the protein FacZ (<u>F</u>irmicute-<u>a</u>ssociated <u>c</u>oordinator of <u>Z</u>-rings). In *S. aureus*, inactivation of the conserved cell division protein GpsB suppressed the division and morphological defects of *facZ* mutants. Additionally, FacZ and GpsB were found to interact directly in a purified system. Thus, FacZ is a novel antagonist of GpsB function with a conserved role in controlling division site placement in *S. aureus* and other Firmicutes.

## MAIN TEXT

The cell envelope of the opportunistic pathogen *S. aureus* is vital for resisting turgor pressure and interfacing with the host. Like most Firmicutes, its surface is composed of a cytoplasmic membrane (CM) surrounded by a thick peptidoglycan (PG) cell wall, with anionic polymers called lipoteichoic acids (LTAs) and wall teichoic acids (WTAs) decorating the respective layers^1^. Because the mechanical integrity conferred by the cell envelope is necessary for survival^2^, the biosynthetic pathways that build it have been effective targets for many of our most successful antibiotic classes, and many strains of *S. aureus* have evolved resistance to cell wall-targeting antibiotics^3^. A deeper understanding of the mechanisms that promote envelope biogenesis therefore promises to uncover new ways to target this essential structure for drug development.

In rod-shaped bacteria, envelope assembly is performed in two phases: elongation and division. Depending on the organism, elongation proceeds via the insertion of new cell wall at dispersed sites throughout the cell cylinder or at the cell poles^4^. This phase is followed by cell division, which in gram-positive bacteria involves the assembly of a multilayered cell wall septum that initially bisects the mother cell. The septum is built by the divisome, a multiprotein PG synthesis machine organized by treadmilling polymers of the tubulin-like FtsZ protein that condense into a dynamic structure called the Z-ring to establish the division site^4^. After the septum is built, enzymes with cell wall cleaving activity called PG hydrolases or autolysins split the septum to promote daughter cell separation and complete the division process^5, 6^.

Spherical *S. aureus* cells do not have as distinct an elongation phase as their rod-shaped cousins^7^. Instead, envelope biogenesis is principally focused at the division site throughout the cell cycle^8^. In many rod-shaped bacteria, a combination of the Min system and nucleoid occlusion define the division site by guiding Z-ring formation to midcell^9–11^. Although *S. aureus* also utilizes a nucleoid occlusion protein^12, 13^, much less is known about how it regulates division-site placement to localize envelope assembly. As a sphere, *S. aureus* must restrict division to a single midcell plane among the theoretically infinite number of such planes available. An additional complexity in *S. aureus* not shared by rod-shaped cells is that its division site rotates through different perpendicular planes in successive cell cycles, giving rise to the characteristic “clustered grape” morphology of cells before completion of daughter cell separation^14^. Geometric and/or structural epigenetic cues have been suggested to aid in coordinating division site rotation, but the underlying mechanism is unclear^15, 16^. Thus, much remains to be learned about how this seemingly simple spherical bacterium controls the biogenesis of its envelope.

To uncover new factors involved in envelope assembly in *S. aureus*, we performed two related comprehensive screens for mutants with altered cell morphology and/or increased envelope permeability. Hits included many genes known to be critical for proper envelope assembly, indicating that the screens worked as intended. The screens also implicated a number of genes of unknown function in envelope assembly, providing a rich dataset for uncovering new insights into the process. We characterized one of these hits, *facZ*, and found that it encodes a conserved regulator of cell division placement.

## RESULTS

### High-throughput screens for S. aureus mutants defective in envelope biogenesis

A high-density transposon mutant library of *S. aureus* was constructed and subjected to two enrichment regimens using a cell sorter (**Fig. 1**). The END (<u>en</u>velope <u>d</u>efective) enrichment sorted for mutants with altered membrane permeability based on increased staining with propidium iodide (PI), a membrane impermeable fluorescent DNA dye (**Fig. 1A**). Although PI staining is typically associated with lethal loss of membrane integrity^17^, we reasoned that we could enrich for mutants experiencing an increased frequency of these defects based on the association of dead, PI-positive cells with viable siblings in unseparated cell clusters. In the related CSD (<u>c</u>ell <u>s</u>eparation or <u>d</u>ivision) enrichment, mutants defective for cell separation or cell division were sorted based on increased light scattering of larger cells or cell clusters^18, 19^ (**Fig. 1B**). To set the sorting cutoffs (gates) for each enrichment, we used a Δ*atl* mutant defective in the major cell separation autolysin^5, 20^ as a control (**Fig. 1C**). As expected, the larger size and shape complexity of unseparated cell clusters of the mutant gave rise to a population of cells causing increased light scattering (Scatter^HI^) compared to a wild-type control. Additionally, we observed the appearance of a significant population of PI-positive (PI^HI^) cells in cultures of the Δ*atl* mutant that was largely absent in wild-type cultures (**Fig. 1C**). With the sorting parameters defined to capture Scatter^HI^ and PI^HI^ cells with similar properties to those of the Δ*atl* mutant, we performed the enrichments on the transposon library (**Fig. 1D**).

**Figure 1.**
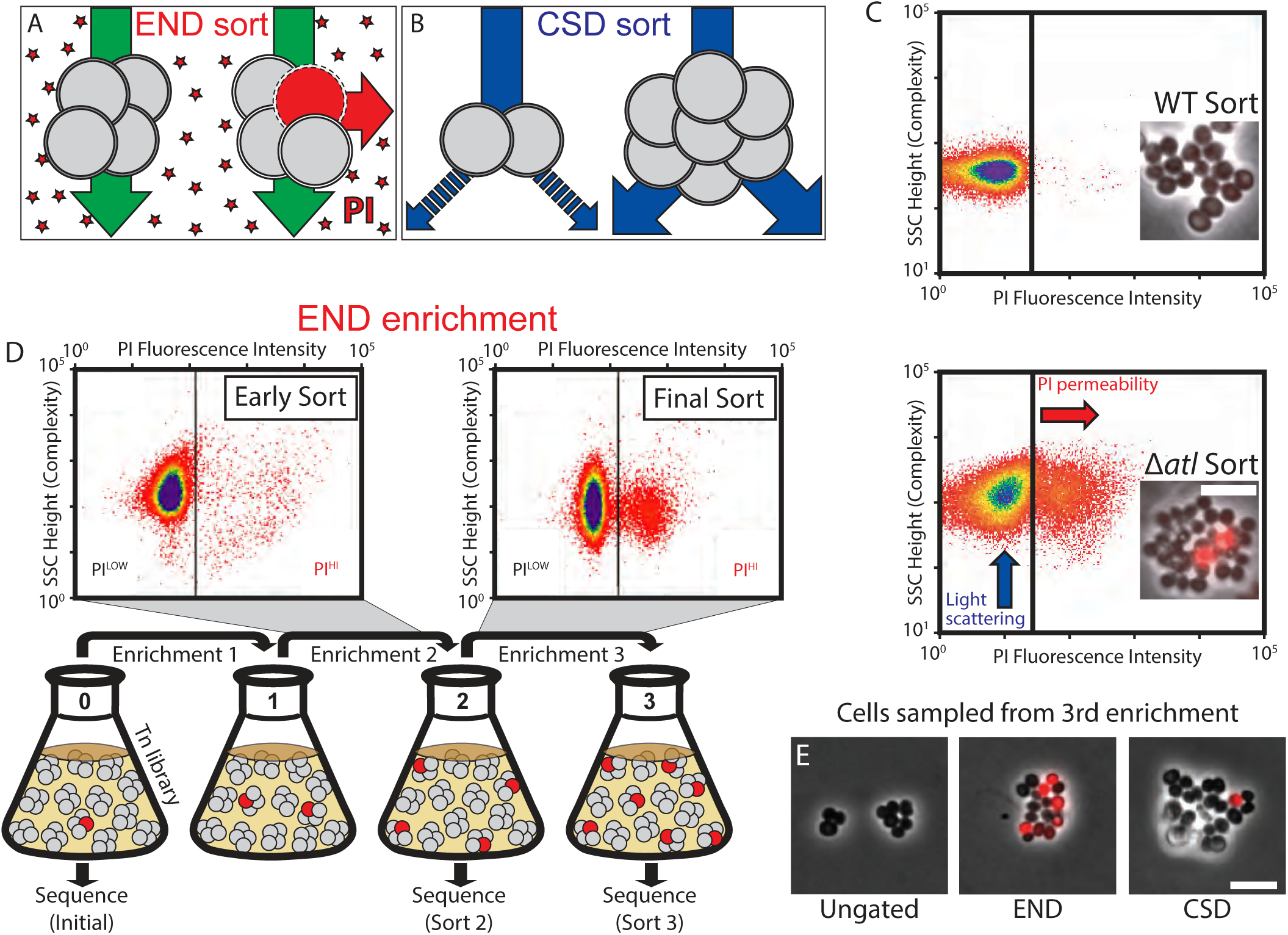
High-throughput FACS-based screens for mutants defective in envelope assembly. A-B) Schematics showing the logic of the END and CSD enrichments. See text for details. C) FACS profiles monitoring light scattering (SSC Height) and propidium iodide (PI) fluorescence for WT and Δ*atl* strains. Inset micrographs show phase contrast images with overlay of PI staining (bar = 4µm). D) Schematic detailing the END enrichment workflow. An analogous CSD enrichment was performed in parallel (not shown). See text for details. Enrichment for PI^HI^ mutants was examined by FACS profiling at each stage (top). E) Representative images of the final cell populations from the END and CSD sorts, as well as an unsorted control, were imaged on 2% agar pads (bar = 4 μm). PI staining (red) was overlaid on the phase contrast image.

Prior to sorting, the library was grown to mid-exponential phase. We then collected cells passing the Scatter^HI^ or PI^HI^ gates (10^6^ cells for each) as well as an equivalent number of cells sorted without a gate for use as a control. The resulting sorted populations were recovered, amplified in growth medium, and sorted again using the same parameters. Three sequential enrichments were performed for each population, maintaining cells in exponential phase throughout the procedure (**Fig. 1D**). Imaging of cells from the populations recovered following the third sort showed that the protocol was effective in enriching for cells with the desired phenotypes (**Fig. 1E**).

To monitor changes in the representation of transposon mutants in the library during enrichment, transposon sequencing (Tn-Seq) was performed on the original unsorted library and the mutant populations following the second and third sorts for Scatter^HI^, PI^HI^, or the ungated control sort. In the initial transposon library, the median gene had 17 unique insertions, and 2,468 of the 2,629 annotated genes (93.9%) had at least one insertion (**Table S1**). To determine which mutants were being enriched by the sorts, the number of insertion reads at each locus from the END or CSD enrichments were compared to those in the ungated control at the same time point (**Table S2**). By the second sort, we observed a subset of genes displaying >4-fold enrichment in insertions relative to the ungated control, and this enrichment became even more pronounced by the third sort (**Fig. S1A-B**). Furthermore, for each enrichment, the increased representation of insertions in a given gene following the second sort was correlated to its enrichment following the third sort (**Fig. S1C-D**). Thus, each enrichment was selective, and multiple rounds of sorting yielded stronger phenotypic enrichment.

The enrichments of END and CSD mutants were moderately correlated, with many of the fifty most-enriched mutant loci for each sorting scheme overlapping (**Fig. S1E, Table S3**). To identify mutants showing the greatest combined enrichment across both sorts, we analyzed the enrichment data using a two-dimensional principle-component analysis (2D PCA) (**Fig. S1F**). A total of 74 candidate ‘hits’ were enriched at least 8-fold compared to unsorted controls along the first principal component, PC1, which serves as a proxy for combined enrichment in both screens (**Table S4**). Among the hits were genes with known roles in cell morphogenesis and cell division like *atl*^5, 20^ and *sagB*^21–23^, with transposon insertions in both genes showing clear enrichment following both the END and CSD sorts (**Fig. 2A, S2A, and Table S5**). Genes of unknown function were also well represented among the list of hits, including *facZ* (*SAOUHSC_01855*), which showed an enrichment in transposon insertions following both sorting methods, and displayed aberrant cell morphology and an increased frequency of PI^HI^ cells in cultures (**Fig. 2, S2A, and Table S5**). Mutants in genes of unknown function SAOUHSC_01975 and SAOUHSC_02383 were also identified as hits, and demonstrated to have defects in cell division and/or envelope permeability (**Fig. S2**). Among all of the hits, we found *facZ* (SAOUHSC_01855) to be particularly interesting because the unusual nature of the membrane and cell wall depositions observed in cells lacking FacZ (**Fig. 2B** and below). We therefore focused on investigating its function.

**Figure 2.**
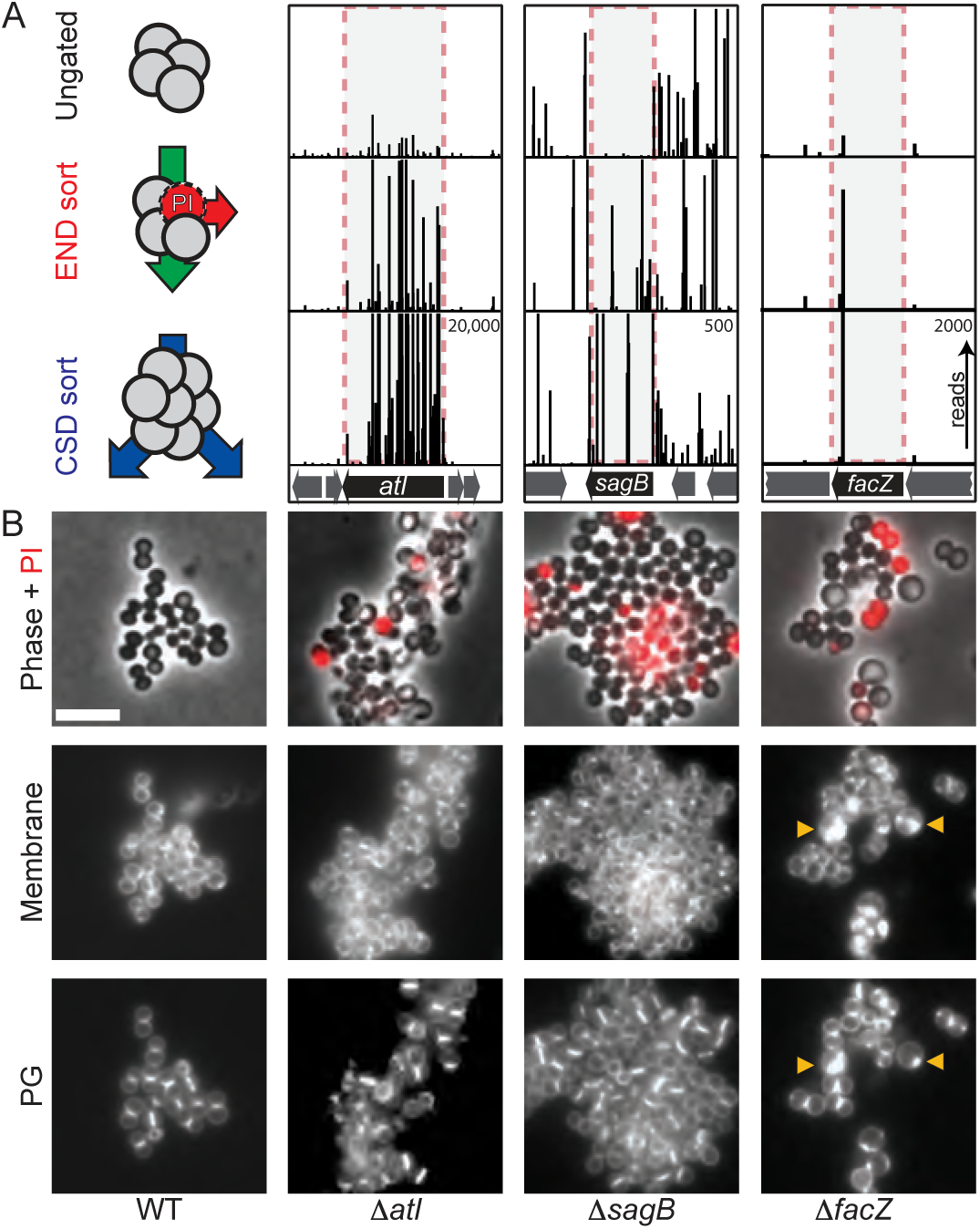
Validation and initial characterization of screening hits. A) Tn-Seq profiles from three genomic loci displaying enrichment of transposon insertions at the completion of the END and CSD sorting protocols relative to the control sort. Each vertical line represents a mapped insertion site, and the height of the line is the number of reads mapping to that site, which reflects the representation of the insertion mutant in the population. Profiles for each locus are scaled separately with the maximum number of reads indicated in the top right corner of the bottom profile. B) Representative images of WT and mutant cells pulse-labeled with sBADA to visualize peptidoglycan (PG) synthesis (bottom row), stained with TMA-DPH to label cell membrane (membrane, middle row), and treated with PI to assess envelope permeability (Phase + PI, top row). Yellow carets highlight membrane and PG synthesis defects. Fluorescence intensity for each channel was scaled identically for all strains to facilitate direct comparison between images (bar = 4 µm).

### facZ mutants have pleiotropic envelope defects

To further characterize the envelope defects of Δ*facZ* cells, we pulse labeled them with the fluorescent D-amino acid (FDAA) sBADA to label sites of nascent cell wall synthesis^24^, followed by treatment with Nile Red and DAPI to visualize the membrane and nucleoid, respectively. The mutant cells were more heterogeneous in size than wild-type cells and displayed strikingly aberrant membrane and PG accumulations that often colocalized (**Fig. 3A-C, S3A**). These accumulations excluded the nucleoid, indicating that they projected into the cytoplasmic space (**Fig. 3D**). Transmission electron microscopy (TEM) also revealed the aberrant accumulation of envelope material in Δ*facZ* mutant cells (**Fig. 3A**), and 3D-SIM super-resolution microscopy of labeled cells confirmed the co-localization of membrane and PG stains (**Fig. S3B**). Furthermore, this analysis revealed that the projections are likely invaginations that are continuous with the peripheral cell envelope (**Fig. S3B and Video 1**). Cells lacking FacZ also displayed aberrant localization of FtsZ-GFP. Many cells had multiple FtsZ structures oriented at oblique angles relative to one another, indicative of problems with division site placement (**Fig. S3C-D**). Expression of a chromosomally integrated copy of *facZ* under control of the anhydrotetracycline (aTc) inducible *tet* promoter (P*tet*) complemented the morphological and growth defects of Δ*facZ* cells, indicating that that the defects were caused by the lack of FacZ production and not effects of the deletion allele on the expression of nearby genes (**Fig. S3D-E**). We conclude that FacZ is critical for normal envelope biogenesis and that its absence results in the formation of spurious invaginations of the envelope that we suspect represent aberrant division attempts.

**Figure 3.**
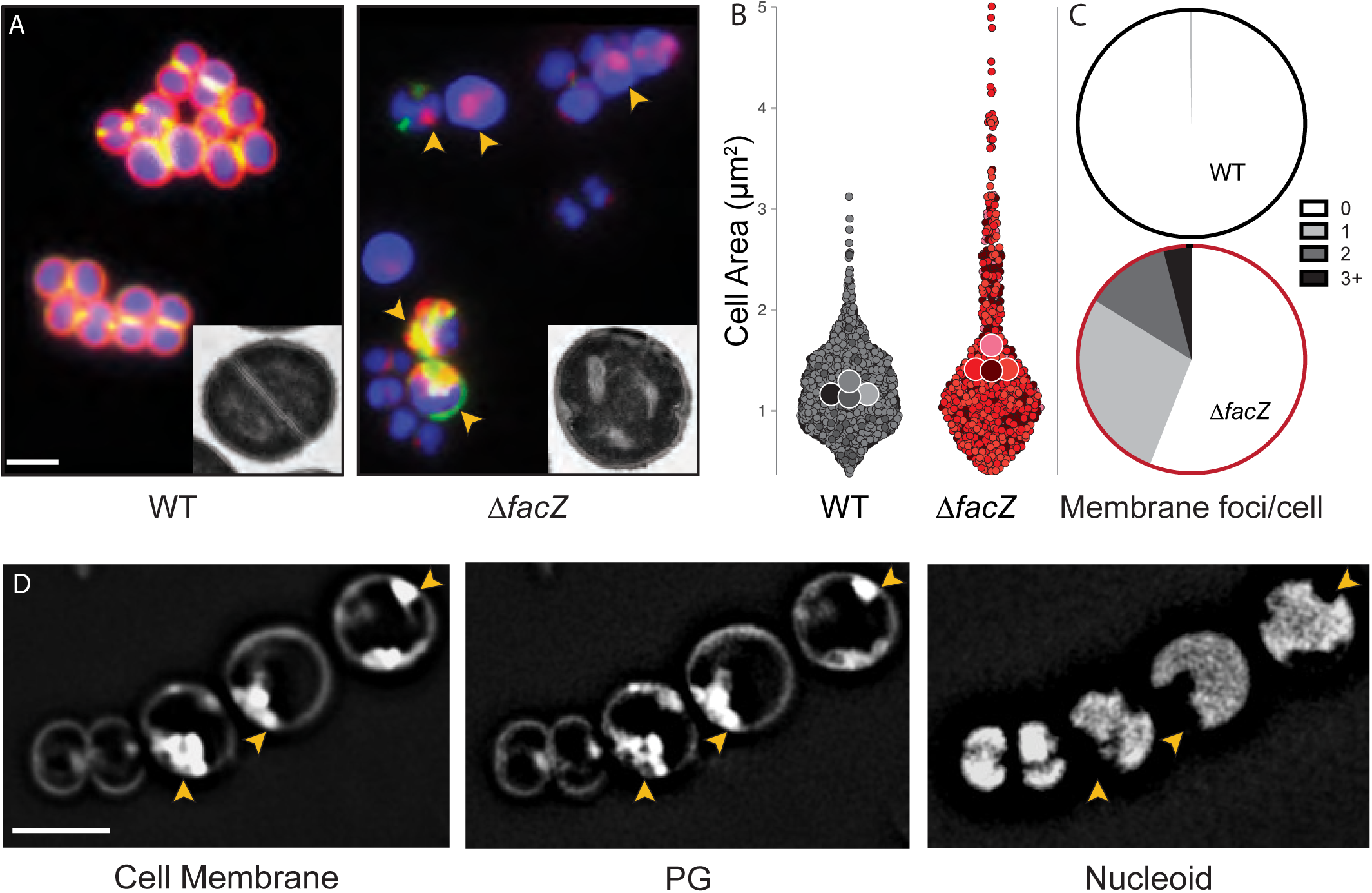
Analysis of morphological defects displayed by Δ*facZ* cells. A) Cells from WT and Δ*facZ* strains were pulse-labeled with sBADA to visualize peptidoglycan synthesis (green), washed three times with PBS to arrest growth and remove unincorporated sBADA, and then labeled for five minutes with the membrane stain Nile Red (red). Cells were imaged on 2% agar pads containing DAPI to visualize the nucleoid (blue). Insets show cells from the same strains imaged by TEM. Yellow carets highlight cells with aberrant membranes and PG synthesis. B) Violin plots showing the cell area of the indicated strains harboring a cytoplasmic fluorescent protein. The results from four biological replicates were pooled, each shaded differently; small circles are individual measurements, and large circles are medians from each replicate (n > 700 cells; see **Fig. S3**). C) Quantification of aberrant membrane foci in TMA-DPH-labeled cells (n > 200 cells; see **Fig. S3A**). D) 3D-SIM microscopy of Δ*facZ* cells stained identically to those in panel (A). Carets highlight the position of aberrant cell wall and membrane accumulations that exclude the nucleoid. Bars = 2 µm.

### FacZ has a conserved role in envelope biogenesis

FacZ is conserved in many Firmicutes, including *B. subtilis* (**Fig. S4**). We previously found that inactivation of the *B. subtilis* homolog of FacZ (*^Bs^*FacZ, formerly called YtxG) resulted in aberrant membrane invaginations and caused defects in sporulation^25^. Our results in *S. aureus* prompted us to reinvestigate the effect of inactivating *^Bs^*FacZ in vegetative *B. subtilis* cells. We confirmed that cells deleted for *^Bs^facZ* display large, aberrant membrane invaginations via Nile Red membrane labeling (**Fig. S5A**). Additionally, we found that *^Bs^*FacZ inactivation caused misplaced FtsZ-GFP structures and led to an increase in cell length, consistent with a division defect (**Fig. S5B-E**). We therefore conclude that FacZ function is likely to be at least partially conserved throughout many Firmicutes.

### Periseptal localization of FacZ in S. aureus

FacZ is an 18 kDa protein predicted to possess an N-terminal transmembrane (TM) helix and a C-terminal cytoplasmic domain composed of a membrane-proximal coiled-coil region (CCR) followed by an intrinsically disordered region (IDR) with a net positive charge (**Fig. S6A-B**). AlphaFold-Multimer predicts that FacZ forms homo-oligomeric assemblies with 3-4 protomers with a well-packed interface. In such a configuration, the complex is predicted to be shaped like an anchor, extending ∼15 nm into the cytoplasm with disordered C-terminal “fingers” that extend from a rigid helical rod (**Fig. S6C**).

To determine the subcellular localization of FacZ in *S. aureus*, we constructed a functional *facZ-mCherry* fusion (**Fig. S6D**) and expressed it from an ectopic locus under control of the P*_tet_* promoter in cells that were also pulse-labeled with the FDAA sBADA to monitor PG synthesis. Consistent with its predicted TM helix, FacZ-mCherry displayed a peripheral membrane localization, outlining all cells in the population (**Fig. 4A, S6D-E**). In addition to the peripheral signal, FacZ-mCherry was enriched at sites of septum formation but appeared to remain confined to the periphery of the septum (the “periseptum”) even at late stages in division (**Fig. 4A, S6D-E**). We quantified this enrichment by measuring two different fluorescence intensity profiles in populations of labeled cells: one through the cell body perpendicular to the septum (orthogonal profile) and the other around the cell periphery (peripheral profile) (**Fig. 4C-D**). In cells in the early stages of division identified by their incomplete PG septa, FacZ-mCherry showed enrichment at the edges of the orthogonal profile, and an enrichment at the periseptum in the peripheral profile where PG labeling was also most intense (**Fig. 4C-D**). These intensity profiles are as expected for a membrane protein and mirror profiles obtained for cells labeled with the membrane dye Nile Red (**Fig. 4E**). However, the FacZ-mCherry localization pattern was strikingly different from PG and membrane profiles in cells at later stages of division where the septum was observed to be complete or nearly so (**Fig. 4C-E**). In these cells, the PG and membrane stains were enriched at midcell in the orthogonal profile as expected, but the FacZ-mCherry signal remained at or near baseline as if it were excluded from the septum (**Fig. 4C-E**). No change was observed in the peripheral profile, as the structure of the periseptum is not expected to change until the final stage of division when cells separate^26^ (**Fig. 4D**). Based on these observations, we conclude that FacZ is enriched at midcell, but unlike typical division proteins, is confined to the periseptal region.

**Figure 4.**
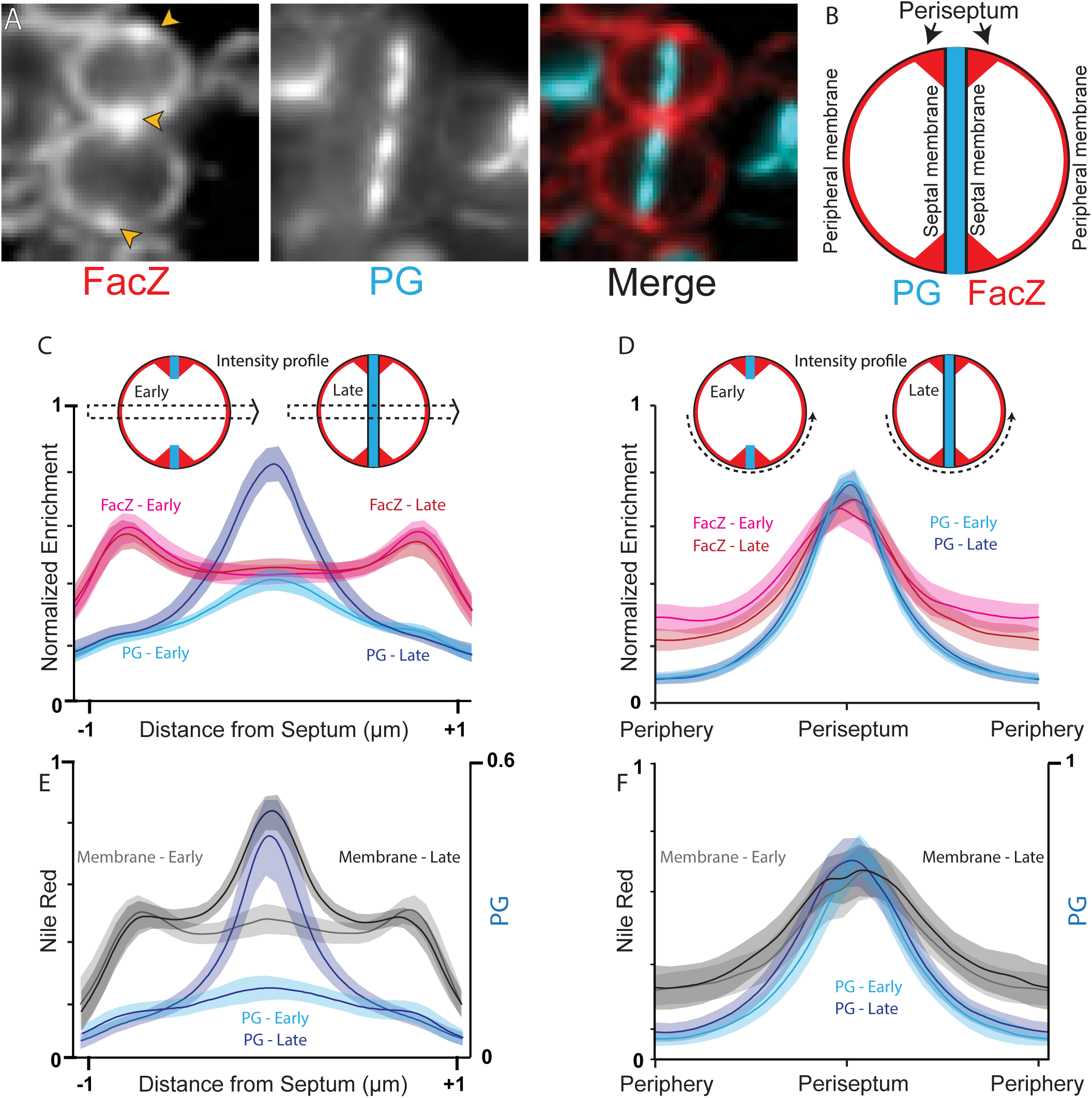
Periseptal localization of FacZ in dividing *S. aureus*. A) Representative fluorescence images of WT cells expressing FacZ-mCherry stained with HADA to label the peptidoglycan (PG). The protein fusion (left) and peptidoglycan (PG) label (center) and a pseudo-colored merged image are shown (FacZ-mCherry in red, HADA in blue). See also **Fig. S6D-E**. B) Schematic depicting the membrane localization of FacZ in a dividing *S. aureus* cell. C) Graphs of mean fluorescence intensity of HADA (light and dark blue) or FacZ-mCherry (light and dark red) collected along lines perpendicular to the septum of cells labeled as in panel (A). Central peaks show intensity at the septum, and peripheral peaks show intensity at the cell periphery. Cells were grouped into early-division (cells with two HADA foci representing an unresolved septum) or late-stage division (cells with a single central HADA focus representing a resolved septum). HADA-labeling is increased in the center of late-stage dividing cells (Welch’s t-test, p < 0.05). Localization of FacZ-mCherry does not change significantly as the septum resolves (Kolmogorov-Smirnov test, p > 0.05). D) Graphs of intensity profile scans measuring fluorescence intensity along peripheral arcs centered on the periseptum for early-and late-stage dividing cells. E-F) Graphs of intensity profile scans of WT cells stained with HADA (light and dark blue) and Nile Red (light and dark grey) imaged and analyzed as in (D) and (E), respectively. Perpendicular intensity profiles (C and E) were normalized from 0-1 for each cell. Peripheral intensity profiles (D and F) were interpolated using MATLAB, and intensity profiles were normalized from 0-1 for each fluorescence channel within each experiment, to facilitate comparison of cells of different size; n ≥ 50 cell segments measured for each condition in each graph.

### A role for FacZ in controlling Z-ring formation

To investigate what aspect of cell division FacZ participates in, we used Tn-Seq to identify synthetic lethal partners. Transposon libraries were generated in wild-type and Δ*facZ* cells, and the insertion profiles were compared (**Table S6**). Notably, insertions in *ezrA*, which encodes a regulator of Z-ring formation^27–,29^, were dramatically depleted in the Δ*facZ* library relative to wild-type (67.6-fold depletion in Δ*facZ*, p < 0.005), suggesting that EzrA is essential in the absence of FacZ (**Fig. 5A**). We confirmed this synthetic lethal relationship by constructing an EzrA depletion strain and demonstrating that EzrA depletion is lethal in a Δ*facZ* background (**Fig. 5B**). Notably, mutants lacking either *ezrA* or *facZ* show increased mislocalization of FtsZ-GFP (**Fig. S3C-D**), and both single mutants are hypersensitive to PC190723 (**Fig. 5C, Fig. S6F**), a chemical inhibitor of cell division that promotes hyper-stabilization of FtsZ filament bundles in Firmicutes^30^. Based on these genetic results, we infer that like EzrA, FacZ plays a role in controlling Z-ring formation and that it may function at the level of FtsZ polymerization.

**Figure 5.**
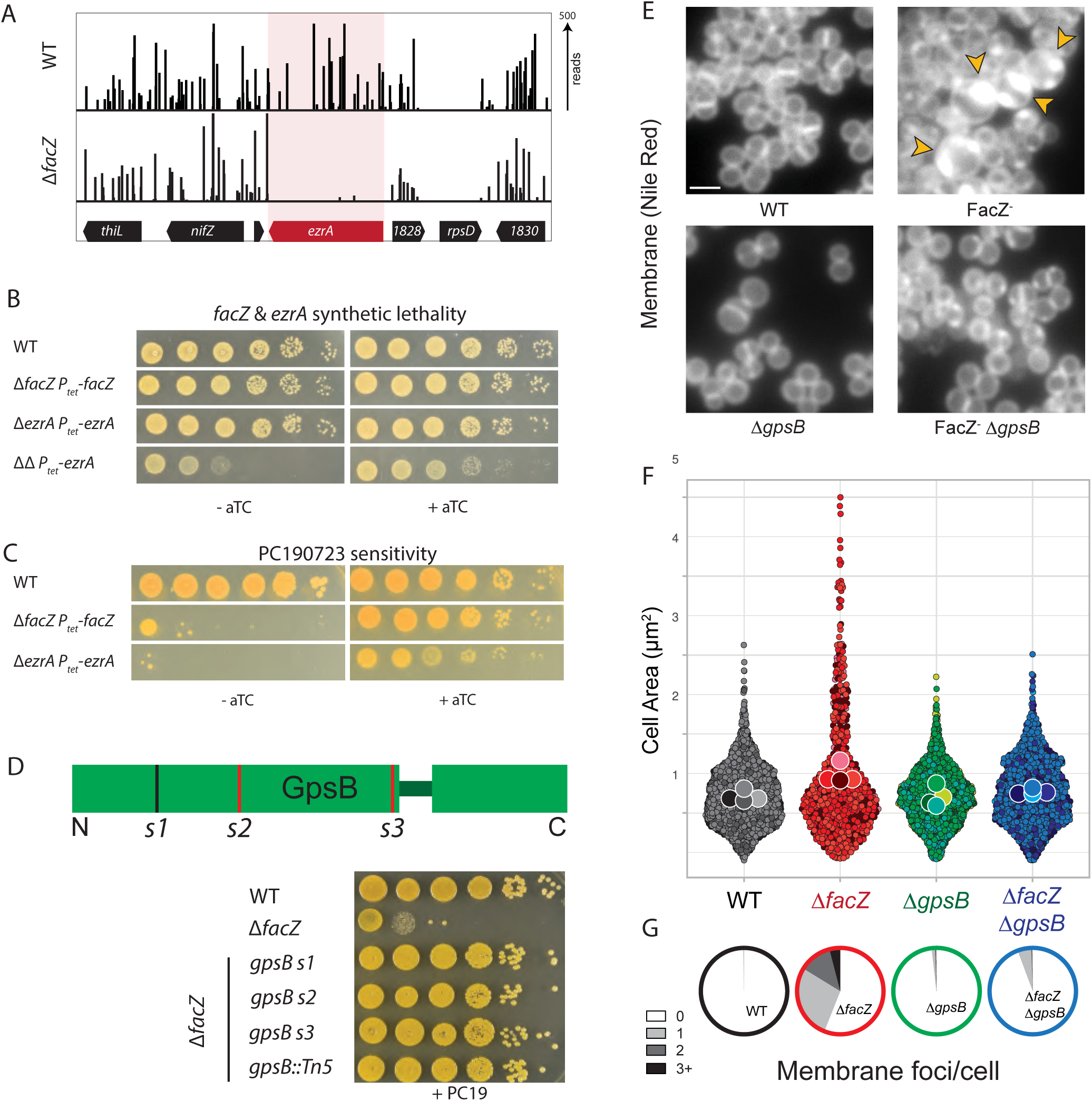
Inactivation of facZ impairs cell division and is rescued by deletion of ΔgpsB. A) Transposon insertion profiles of the *ΔfacZ* locus in strains aTB015 [WT] and aTB259 [Δ*facZ*]. B) Spot titers of cultures of aTB003 [WT], aTB372 [Δ*facZ* P*tet-facZ*], aTP481 [Δ*ezrA Ptet-ezrA*], and aTB378 [Δ*facZ ΔezrA Ptet-facZ*]. Cells were normalized to OD600 = 1.0, serially diluted, and spotted on TSA agar with or without aTC inducer (50 ng/mL). C) Spot titers of aTB003 [WT], aTB372 [Δ*ΔfacZ Ptet-facZ*] and aTP481 [Δ*ezrA* P*tet*-*ezrA*] as in panel (B) except that the plates contained PC190723 (100 ng/mL). D) Top: Diagram showing the location of mutations in *gpsB* that suppress the PC190723 sensitivity of a Δ*facZ* mutant. Suppressor mutations are mapped onto a diagram of the two folded domains of GpsB (lines indicate position of mutations, with red lines indicating mutations generating premature stop codons). Bottom: Cultures of strains aTB003 [WT], aTB251 [Δ*facZ*], aTB453 [Δ*facZ gpsB*-s1], aTB476 [Δ*facZ gpsB*-s2], aTB478 [Δ*facZ gpsB*-s3], and aTB497 [Δ*facZ gpsB*::*Tn*] were OD-normalized and spotted on TSA supplemented with PC190723 (100 ng/ml) as in panel (B). E) Representative fluorescence images of aTB003 [WT], aTB372 [Δ*facZ* P*tet*-*facZ*], aTB525 [Δ*gpsB*], and aTB540 [Δ*gpsB* Δ*facZ* P*tet*-*facZ*]. Strains were grown to mid-log phase without induction of *facZ*, and membranes were stained with Nile Red (see also **Fig. S7**). Yellow carets highlight membrane defects. E-F) Cultures of aTB521 [WT], aTB527 [Δ*facZ*], aTB529 [Δ*gpsB*], and aTB542 [Δ*facZ* Δ*gpsB*] constitutively expressing cytoplasmic RFP from pKK30 were labeled with TMA-DPH in mid-log phase and imaged on 2% agarose pads. (F) Violin plots showing cell area of indicated strains based on cytoplasmic fluorescence (n > 700 cells). (G) Aberrant membrane foci were quantified (n > 200 cells).

### FacZ restricts GpsB to midcell

To further explore the role of FacZ in division, we selected for suppressor mutations that overcome the hypersensitivity of Δ*facZ* cells to PC190723. Whole genome sequencing of the isolated suppressors identified three unique mutations in *gpsB*, a gene encoding a conserved Firmicute-associated protein with a role in envelope biogenesis^31^ that has been shown to interact with FtsZ in *S. aureus*^32, 33^. The lesions included an in-frame duplication of residues 21-24 and two deletions causing premature stop codons at residues 34 and 66, all of which are likely to disrupt GpsB function (**Fig. 5D**). Although GpsB has been proposed to be essential in *S. aureus*, previous Tn-seq analyses from our labs and others have identified Tn mutants throughout *gpsB* in several strains of *S. aureus*^12, 34^, and the NTML collection of nonessential *S. aureus* genes contains a Tn mutant in *gpsB*^35^. Accordingly, we constructed a kanamycin-marked Δ*gpsB* (Δ*gpsB::kan^R^*) strain. Both this marker and the Tn-inactivated allele from the NTML collection were readily transduced between strains, indicating that *gpsB* is dispensable for growth in our conditions. As expected from the suppressor analysis, deletion of *gpsB* alleviated the PC190723 hypersensitivity of Δ*facZ* cells (**Fig. 5D**). Importantly, GpsB inactivation also largely suppressed the cell size and membrane invagination defects caused by deletion of *facZ* (**Fig. 5E-G and S7**).

Localization of FacZ-mCherry and mNeon-GpsB fusions indicate that both proteins are present at the periseptal region in cells at early stages of division (**Fig. 6A-B**). However, FacZ-mCherry remains at the periseptum in later stages of division while mNeon-GpsB localizes throughout the septum (**Fig. 6A-B**). Notably, the localization of mNeon-GpsB was altered upon inactivation of FacZ, with the fusion enriching at many sites throughout the cell coincident with areas of aberrant membrane invagination (**Fig. 6C**).

**Figure 6.**
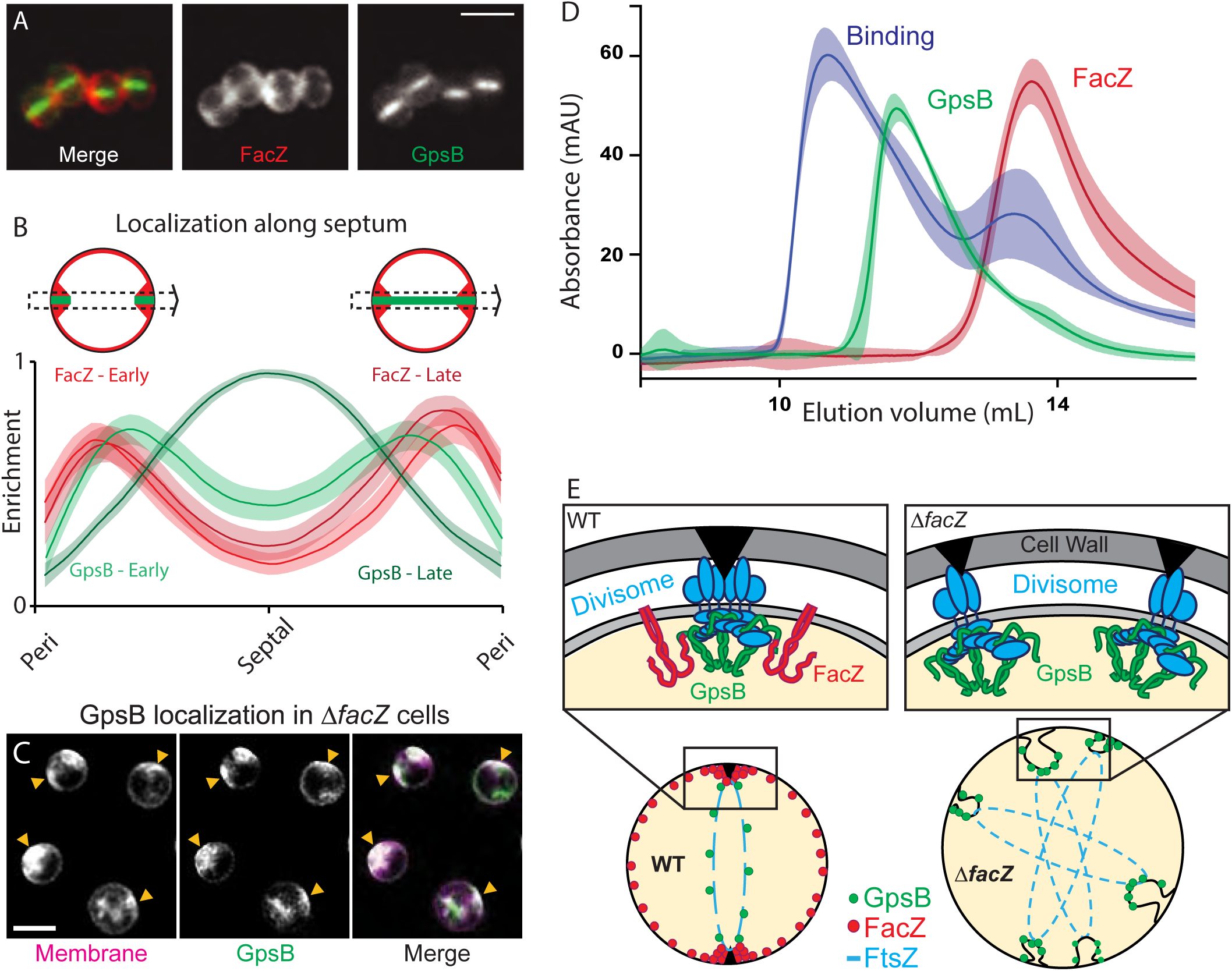
FacZ interacts with GpsB and influences its localization. A) Representative images of aTB517 [Δ*facZ* Δ*gpsB Ptet-gpsB-mNeon Pspac-facZ-mCherry*] grown in the presence of aTC (50 ng/mL) and IPTG (25 ng/mL) and imaged by fluorescence microscopy. B) Graphs showing the fluorescence intensity of FacZ-mCherry (light and dark red) and GpsB-mNeon (light and dark green) along lines perpendicular to the septum. Cells were separated into two groups based on the GpsB-mNeon signal: cells with discontinuous GpsB foci were considered early-division cells (top left) whereas cells with a continuous GpsB-mNeon band at the septum were considered late-division cells (top right). (n ≥ 50 cells for each group). C) Representative deconvolved fluorescence images of GpsB-mNeon localization in Δ*facZ* cells (aTB519) labeled with Nile Red. D) Gel filtration profiles of GpsB (1-75) alone (green), SUMO-3x-FacZ (127-146) alone (red), or a mixture of the two proteins (binding, blue). Elution profiles represent the average and standard deviation of 6 runs. E) Schematic model of FacZ function within the divisome. Cell division is properly localized when FacZ is functional (left). In the absence of FacZ, GpsB is unregulated such that cell constriction is initiated at many sites (right). See text for details.

These findings suggest that FacZ functions by constraining GspB localization and activity to a single site to promote normal division.

To determine if FacZ directly interfaces with GpsB to control its function, we tested whether the two proteins interact in a purified system. Previous crystal structures show that the N-terminal domain of GpsB forms a dimer that binds to small peptides of *S. aureus* FtsZ and PBP4 bearing a (S/T/N)RxxR(R/K) motif^31, 36^. We reasoned that FacZ might interact with GpsB by a similar mechanism and found that it contains a canonical *Staphylococcal* GpsB-binding motif at residues 134-139 (NRHYRR). We purified a SUMO-tag fusion bearing a 3x concatenation of residues 127-145 of FacZ (FacZ_127-145_). Incubation of this fusion with the N-terminal domain of GpsB (GpsB_1-75_) resulted in a shift in the elution profile of the two proteins by size exclusion chromatography, indicating that the two proteins interact (**Fig. 6D-E**). These data support a model in which FacZ functions in cell division by directly interacting with GpsB and antagonizing its activity to prevent spurious Z-ring formation and aberrant envelope invaginations (**Fig. 6F**).

## DISCUSSION

Our genetic screens for envelope biogenesis factors identified FacZ as a new regulator of division-site placement in *S. aureus*. Cells lacking FacZ have aberrant divisome structures, increased cell size, and invaginations of the cell envelope, suggesting that in the absence of FacZ, FtsZ forms spurious divisome complexes that initiate constriction but fail to complete cytokinesis. Importantly, the morphological defects observed in a Δ*facZ* mutant are largely suppressed when *gpsB* is deleted. Furthermore, GpsB localizes to the aberrant constrictions formed in the Δ*facZ* mutant, and FacZ harbors a GpsB-interacting sequence that promotes its direct association with GpsB *in vitro*. Thus, our data support a model in which FacZ functions by directly interacting with and constraining the activity of GpsB to prevent aberrant division events.

GpsB was previously shown to interact directly with FtsZ and to promote lateral interactions between FtsZ filaments^32^. It has also been shown to interact with several factors involved in cell envelope biogenesis, including PG synthases and enzymes involved in WTA biogenesis^33, 36^. Thus, GpsB has been proposed to function in the stabilization of Z-rings at the onset of cell division and in the recruitment of enzymes that participate in septum synthesis to the division site. The mechanism by which FacZ restrains GpsB function is currently unknown, but we envision two possible scenarios, both of which rely on the ability of FacZ to directly or indirectly localize to sites where membrane invagination is being initiated by full or partial divisome complexes. Such an activity is supported by the enrichment and retention of FacZ-mCherry at the periseptum throughout the division process, which suggests a preference for sites of strong positive Gaussian curvature such as those formed at the junction of the septal and peripheral membrane during invagination.

In the first potential mechanism, FacZ localizes to membrane invaginations at sites outside of the preferred division site, which is defined by other systems like nucleoid occlusion. It then blocks maturation of these early-stage constrictions by inhibiting the ability of GpsB to interact with FtsZ and/or other divisome components at these ectopic sites. Alternatively, FacZ could function to reinforce the preferred division site by enhancing the ability of GpsB to promote Z-ring formation and divisome maturation. In this second scenario, the absence of FacZ would reduce the difference in divisome-forming potential between preferred and non-preferred sites and thus promote aberrant envelope synthesis. In both cases, we propose that inactivation of GpsB restores normal division by making Z-ring formation and divisome maturation more reliant on signals from other division localization systems that define the preferred site. Further work will be required to differentiate between these and other possible models for FacZ function, and to determine why cells have the FacZ-GpsB system if cell division can proceed relatively normally in the absence of both proteins. Like many other division-site placement systems, FacZ and GpsB are nonessential in laboratory conditions, but their broad conservation within the Firmicutes suggests they provide a strong selective advantage for growth in the wild.

FacZ was just one of many previously uncharacterized proteins that our genetic enrichments have implicated as playing important roles in envelope biogenesis and cell division in *S. aureus*. The dataset we generated should therefore be a valuable resource for uncovering additional new insights into the mechanisms used by this pathogen to control the construction of its surface. These discoveries, in turn, will enable future efforts aimed at targeting this essential process for the discovery of novel antibiotics to overcome resistance.

## Methods

### General Methods

All cells were streaked for single colonies on 1.5% agar plates of the appropriate culture medium prior to experiments. *S*. *aureus* strains were grown in tryptic soy broth (TSB) at 30 °C or 37 °C with aeration, unless otherwise indicated. Cultures were supplemented where necessary with antibiotics at the following concentrations: erythromycin (Erm 5 μg/ml for chromosomal insertions and 10 μg/ml for plasmids), chloramphenicol (Cm 5 μg/ml), spectinomycin (Spec 200 μg/ml), kanamycin (Kan 25 μg/ml), and PC190723 (0.2 μg/ml); and induced where necessary with isopropyl ß-d-thiogalactopyranoside (IPTG, 25 or 50 μM), or anhydrotetracycline (aTc, 50 ng/ml unless otherwise indicated). *B*. *subtilis* strains were derived from PY79. Cells were grown in Luria Broth (LB) or Casein Hydrolysate (CH) medium at 37 °C with aeration, unless otherwise indicated. When necessary, media were supplemented with tetracycline (10 μg/ml), Spec (100 μg/ml), Kan (10 μg/ml), Cm (5 μg/ml), xylose (0.5% w/v or IPTG (0.5 mM). *E. coli* strains were grown in LB at 37 °C with aeration and supplemented with ampicillin (100 μg/ml). Experiments were conducted on mid-log phase cells (OD_600_ ∼ 0.4) unless noted.

### Strain Construction

All *S. aureus* deletions were generated in strain RN4220 [WT] by homologous recombination as previously described^12^, with the exception of Δ*gpsB::kanR* (TAS201), which was generated via a recombineering system modified from Datsenko & Wanner^37^, the details of which will be the subject of a separate publication. Complementation strains were initially made in RN4220 via integration of pTP63 into a phage attachment site on the chromosome of strain aTB033 (RN4220 *pTP044*)^12^, or from the multicopy plasmid pLOW^38^ in RN4220. Following initial strain construction, markers were transduced by Φ11 or Φ85 into clean genetic backgrounds. All *B. subtilis* strains were generated using the one-step competence method unless indicated otherwise. *E. coli* strains used to passage plasmids for *B. subtilis* or *S. aureus* were generated by electroporation or chemical transformation into DH5α cells, while *E. coli* strains used for protein purification were produced by chemical transformation of plasmids into BL21 (DE3). Plasmids were constructed by restriction digest and ligation, or isothermal assembly (ITA). Lists of primers, strains, plasmids, and descriptions of their construction can be found in Supplemental Materials (see Tables S7-S10).

### Transposon insertion library construction and sequencing

Transposon library construction and sequencing was based on Pang *et al*^12^. Following isolation of cell pellets by centrifugation, cells were lysed with lysostaphin, then treated with RNAse A (ProMega) and Proteinase K (NEB). Genomic DNA was sonicated (Q800R2 QSONICA) to 200–600 bp fragments, then PolyC-tails were added by incubating fragments with a 20:1 mixture of dCTP (Thermo Scientific) to ddCTP (Affymetrix) and TdT enzyme in TdT buffer (ProMega). This DNA was then subject to an initial round of PCR amplification with primers oTB535 and oTB536 with the Easy A cloning kit (Agilent), digested with NotI (NEB) to remove contaminating genomic DNA, and subjected to a second PCR amplification with primer oTB537 (an Illumina sequencing primer) and a barcoding primer (unique for each sample to allow for demultiplexing of reads) with the Easy A cloning kit (Agilent). A 200–400 bp product was gel-purified and sequenced on an Illumina NextSeq Platform (in lab) or an Illumina HiSeq 2500 platform (TUCF Genomics Facility, Tufts University). Following sequencing, reads were mapped to reference genome NC_007795 and statistically analyzed as previously described^12^. Raw data will be made accessible through NCBI’s Sequence Read Archive upon final publication.

### Suppressor analysis and whole-genome sequencing

*S. aureus* Δ*facZ* was plated onto restrictive conditions (TSA supplemented with 0.1 μg/mL PC190723), and spontaneous suppressors were isolated and cultured overnight, along with wildtype and the parental Δ*facZ* mutant. Cell pellets were collected by centrifugation, then lysed with 20 μg/ml lysostaphin in PBS at 37 °C for 30 minutes. DNA libraries were prepared using the Illumina DNA Prep kit and IDT 10bp UDI indices, and sequenced on an Illumina NextSeq 2000, producing 2×151bp reads. Demultiplexing, quality control, and adapter trimming was performed with bcl2fastq2 (v2.20.0.422). Variant calling was performed using Breseq (version 0.35.4)^39^. Single nucleotide polymorphisms (SNP) and deletions were identified by comparing the sequence of the suppressors to the parental strains. Raw data will be made accessible through NCBI’s Sequence Read Archive upon final publication.

### Phenotypic enrichment by FACS

FACS was performed with an Astrios FACS (Beckman Coulter) sterilized with bleach and washed with filtered sterile ddH_2_O before and after each sort. For both END and CSD sorts, wild-type *S. aureus* was compared to a Δ*atl* mutant to set phenotypic gates that would exclude normal cells and enrich for mutants with similar phenotype to the Δ*atl* strain. The Tn-library of *S. aureus* RN4220 (described above) was then sorted using these gates alongside a control that was passed through the FACS but without any sorting gates set. The first 10^6^ cells to pass through the FACS that satisfied the sorting parameters were used to inoculate a 500 ml flask of TSB at 30°C with aeration. Each population was then subjected to iterated rounds of sorting using the same gates between rounds. Cells were kept in exponential phase throughout sorting, which took place over roughly 36 hours. Aliquots from each sort were removed for immediate imaging and frozen in glycerol at −80°C for Tn-seq analysis.

### Statistical comparison of Tn-seq data sets

Following sorting, sequencing, and mapping, the “read ratio” of each genetic locus was calculated by dividing the number of reads at a given locus in that experimental sort by the number of reads at the same locus in its relevant control population:

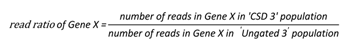

Because each dataset contained a different number of reads, these read ratios were not comparable between them. For example, if an experimentally sorted dataset contained more reads than the control ungated dataset, the mean read ratio for that sort would be greater than one, and this delta would be different for every dataset. To allow comparison between datasets (e.g., tracking the change in enrichment of Gene X from ‘CSD sort 2’ to ‘CSD sort 3’), the “relative enrichment” of each gene was determined by subtracting the difference between the ‘global’ mean read ratio for each population from the ‘specific’ read ratio at each locus:

> *relative enrichment of Gene X = read ratio of Gene X - mean read ratio for relevant sort*

For example, if a given gene had a 2-fold increase in number of reads in a sorting round, but there were twice as many total reads in that sort than in its ungated control, then its relative enrichment would be 2 – 2 = 0 (zero). These normalized (centered) datasets were used for all comparisons between Tn-Seq analyses. Comparison between different rounds of sorting (e.g., CSD sort 2 vs CSD sort 3) was performed by linear regression analysis. Comparison between the CSD and END sorts (e.g., final CSD enrichment vs final END enrichment) was performed by 2D-PCA. In this analytical reference frame, enrichment along PC1 between rounds of sorting serves as a proxy for joint enrichment in both sorts.

### Fluorescence Microscopy

Cells were collected by centrifugation at 3,300 xg for 2 min, then immobilized on PBS or M9 agarose pads (2% wt/vol). Cell walls were stained with FDAAs (HADA or sBADA) (Tocris) at 100 μM. Cell membranes were stained with 1 μg/ml Nile Red (ThermoFisher). DNA was stained with DAPI (Molecular Probes) at 2 μg/ml, or with Propidium Iodide (Molecular Probes) at 5 μM. Standard-resolution fluorescence microscopy was performed using a Nikon Eclipse Ti2 or Ti inverted microscope with a Nikon CFI Plan Apo VC 100X objective lens. 3D-SIM images were acquired with a Nikon Ti2 inverted microscope equipped with an N-SIM Spatial Light Modulator, a Physik Instrument Piezo Z motor, Nikon Laser Illuminators, and captured with a Dual Hamamatsu Orca Flash 4.0 camera using Nikon Elements 5.11 acquisition software.

### Quantitative Image Analysis

Image quantification was performed with MicrobeJ^40^ in ImageJ. Cell identification, segmentation, and morphological assessment was performed on cytoplasmic fluorescence intensity signal. For *B. subtilis*, settings were as previously described^41^. For *S. aureus*, input settings were as follows: background = dark; thresholding = 11; area = 0.4-2.5; options = clump (0.1); all other settings = default. Parameters were kept the same between different strains/conditions within each experiment. All morphology assessments were fully automated with the following exceptions: categorization of cell cycle phase (early vs. late division), number of membrane foci/cell, aberrant vs. normal division, and angle of division plane were manually determined. Furthermore, poorly-separated cells that were difficult to segment or obviously mis-identified by MicrobeJ were manually removed from analysis. Fluorescence intensity scans (see **Fig. 4** and **Fig. 6**) were made from user-defined regions of interest. Because fluorescence intensity measurements have arbitrary units, measurements were normalized from 0-1 (either within each ROI, or within all ROIs measuring the same signal in a given experiment) to allow comparison and graphical display of relative intensity shifts. For measurements of different lengths, measurements were interpolated to facilitate direct comparison between measurements using a custom MATLAB script, which will be made available upon final publication. Fluorescence images were adjusted for display in FIJI and Adobe Illustrator, and images shown side-by-side for comparison were collected and analyzed with identical parameters.

### Electron Microscopy

SEM was performed based on Pang *et al*^12^. Briefly, cells were pelleted and fixed overnight at 4°C in a mixture of 1.25% formaldehyde, 2.5 % glutaraldehyde, and 0.03% picric acid in 0.1 M sodium cacodylate buffer, pH 7.4. The fixed tissues were washed with 0.1 M sodium cacodylate buffer and post-fixed with 1% osmium tetroxide/1.5% potassium ferrocyanide (in H2O) for 2 hours. Samples were then washed in a maleate buffer and post fixed in 1% uranyl acetate in maleate buffer for 1 hour. Samples were then rinsed in ddH20 and dehydrated through a series of ethanol solutions (50%, 70%, 95%, (2x)100%) for 15 minutes each. Dehydrated tissues were put in propylene oxide for 5 minutes before they were infiltrated in epon mixed 1:1 with propylene oxide overnight at 4°C. Samples were polymerized in a 60°C oven in epon resin for 48 hours. They were then sectioned into 80-nm thin sections and imaged on a JEOL 1200EX Transmission Electron Microscope. Images were recorded with an AMT 2k CCD camera.

### Protein purification

A DNA sequence encoding a 3x concatenation of FacZ127-145 separated by GSAG linkers was gene-synthesized and cloned into a vector with an N-terminal His-SUMO tag, with the sequence:

EIADKWQNRHYRRGSANYKA<u>GSAG</u>EIADKWQNRHYRRGSANYKA<u>GSAG</u>EIADKWQNRHYRRGSANYKA (linker sequence underlined). Amino acids 1-75 of *S. aureus* GpsB were cloned into a vector bearing an N-terminal His-SUMO tag. His-SUMO-FacZ127-145 and His-SUMO-GpsB1-75 were purified separately using the same protocol. A 1:100 dilution of an overnight culture of BL21 (DE3) cells carrying expression plasmid was inoculated into LB Kan (50 µg/mL) (1L), grown to OD600 = 0.6 at 37°C, and induced with 0.5 mM IPTG at 18°C for 18 hours. All following steps were performed on ice or at 4°C. Cells were collected by centrifugation and lysed via sonication in 40 mM Tris pH 8, 500 mM NaCl, 10% glycerol, 10 mM imidazole, supplemented with 1 mM DTT, 3 mM PMSF, and 1mM benzamidine. Lysates were clarified via centrifugation and applied to 4 mL of pre-equilibrated Ni^+^-NTA resin (GoldBio) in a gravity flow column, washed extensively, and eluted in 300 mM imidazole. For GpsB1-75, the SUMO tag was cleaved with SUMO protease, followed by re-application to Ni^+^-NTA resin to remove uncleaved protein and His-SUMO tag. Proteins were dialyzed against a buffer containing 20 mM Tris pH 8, 500 mM NaCl, 10% glycerol, 1 mM DTT for >24 hours, concentrated to >10 mg/mL, and frozen at −80°C.

### GpsB and FacZ gel-filtration analysis

His-SUMO-FacZ127-145 and GpsB1-75 were thawed from frozen aliquots and separately buffer-exchanged into an identical buffer (20 mM HEPES, pH 7.6, 500 mM NaCl, 1 mM DTT) using PD10 de-salting columns (GE Healthcare). Gel filtration samples were made by mixing GpsB1-75 at 90 μM with SUMO-FacZ127-145 at 30 μM in 500 μL reactions. Concentrated protein stocks were diluted with HEPES buffer lacking salt to reduce the salt concentration to a final concentration between 150 mM and 200 mM. Samples were fractionated on a Superdex75 Increase 10/300 gel filtration column (Cytivia) equilibrated in a buffer containing 20 mM HEPES, pH 7.6, 100 mM NaCl, 1 mM DTT (flow rate - 0.5 mL/minute).

### FacZ conservation analysis

FacZ was mapped to representative dendrograms of bacteria using AnnoTree version 1^42^. Multiple sequence alignments were carried out using Clustal Omega version 1.2.4^43^.

### FacZ structural predictions

Simple assessment of FacZ topology was performed with Protter^44^. Characterization of the coiled-coil region was performed with the COILS server^45^, and identification of C-terminal IDR was performed with IUPRED2A^46^. Homo-oligomeric assembly predictions were made using AlphaFold Multimer v2.1.2^47^, based upon standard output metrics (overall pLDDT score, pTM score, ipTM score).

### Spot titers

Overnight cultures of *S. aureus* were normalized by optical density (OD600 = 1.0) and subject to 10-fold serial dilution. A 3-µl volume of each dilution was spotted onto TSA agar supplemented with antibiotics or inducers when necessary. Plates were incubated at 30°C or 37°C overnight and imaged the next day.

### Statistics and reproducibility

Unless otherwise noted, experiments were carried out in at least biological triplicate, and images shown are representative of multiple experiments. Graphs either display the results of representative experiments with error bars addressing uncertainty within an experiment, or they show the median/mean of multiple experiments with error bars addressing the uncertainty between experiments, as indicated in the figure legends. Specifically, all widefield microscopy analyses were carried out in at least biological triplicate, and many cells from multiple fields of view were analyzed. All spot titers were carried out in at least biological triplicate. *In vitro* binding assays were performed in triplicate using proteins from two independent purifications (6 total replicates) for all three conditions assayed (FacZ alone, GpsB alone, and FacZ+GpsB). Electron microscopy and 3D-SIM imaging were not reproduced, as they primarily serve to improve image resolution for display purposes, and in all cases confirm the findings of successfully replicated widefield experiments. Similarly, genetic screens were not directly replicated, but all hits were confirmed by genetic analysis, and phenotypes were confirmed and replicated at least three times. In general, p-values were derived from Student’s t-tests when comparing reasonably normal distributions of nearly-equal variance, Welch’s t-tests when variance was not similar, and Kolmogorov-Smirnov tests when distributions were non-normal.

## Author Contributions

Project conception and management was performed by TMB, DZR, and TGB. RWB designed, performed, and analyzed the structural predictions of FacZ and the *in vitro* study of the FacZ-GpsB interaction in consultation with TMB. All other experiments were designed, performed, and analyzed by TMB under guidance of DZR and TGB. AM, TAS, and SW contributed unpublished strains and genetic resources. Manuscript written by TMB, RWB, DZR, and TGB.

## Acknowledgments

We thank the Bernhardt and Rudner labs for support and discussion, especially Andrea Vettiger, Jeremy D. Amon, Josué Flores-Kim, Elayne M. Fivenson, Katherine E. Hummels; the Walker lab for *S. aureus* expertise, especially Julia E. Page and Madeleine C. Stone; Camilla Henriksen for helpful discussion; Joel W. Sher for data analysis resources; and Siân V. Owen for advice and support. Fluorescence microscopy was performed at the Microscopy Resources on the North Quad (MicRoN) core at Harvard Medical School; electron microscopy imaging was performed with HMS Electron Microscopy Facility. TMB is a jointly mentored postdoctoral fellow bridging work in both the Bernhardt and Rudner labs and was funded in part by the Ruth L. Kirschstein Postdoctoral Individual National Research Service Award from the National Institutes of Health (NIH-NIAID, F32AI36431). This work was also supported by the Howard Hughes Medical Institute (TGB) and the National Institutes of Health grants R01AI083365 (TGB), R01GM127399 and R01GM086466 (DZR), R01AI139083 (TGB and DZR), and U19 AI158028 (TGB, DZR, and SW).

## References

1 Silhavy, T. J., Kahne, D. & Walker, S. The bacterial cell envelope. Cold Spring Harb Perspect Biol 2, a000414, doi:10.1101/cshperspect.a000414 (2010).

2 Koch, A. L. Growth and Form of the Bacterial Cell Wall. American Scientist 78, 327–341 (1990).

3 Loomba, P. S., Taneja, J. & Mishra, B. Methicillin and Vancomycin Resistant S. aureus in Hospitalized Patients. J Glob Infect Dis 2, 275–283, doi:10.4103/0974-777x.68535 (2010).

4 Rohs, P. D. A. & Bernhardt, T. G. Growth and Division of the Peptidoglycan Matrix. Annual Review of Microbiology 75, 315–336, doi:10.1146/annurev-micro-020518-120056 (2021).

5 Oshida, T. et al. A Staphylococcus aureus autolysin that has an N-acetylmuramoyl-L-alanine amidase domain and an endo-B-N-acetylglucosaminidase domain: Cloning, sequence analysis, and characterization. Proceedings of the National Academy of Sciences 92, 285–289 (1994).

6 Vollmer, W., Joris, B., Charlier, P. & Foster, S. Bacterial peptidoglycan (murein) hydrolases. FEMS Microbiology Reviews 32, 259–286, doi:10.1111/j.1574-6976.2007.00099.x (2008).

7 Lund, V. A. et al. Molecular coordination of Staphylococcus aureus cell division. eLife 7, e32057, doi:10.7554/eLife.32057 (2018).

8 Pinho, M. G. & Errington, J. Dispersed mode of Staphylococcus aureus cell wall synthesis in the absence of the division machinery. Molecular Microbiology 50, 871–881, doi:https://doi.org/10.1046/j.1365-2958.2003.03719.x (2003).

9 Wu, L. J. & Errington, J. Coordination of Cell Division and Chromosome Segregation by a Nucleoid Occlusion Protein in Bacillus subtilis. Cell 117, 915–925, doi:10.1016/j.cell.2004.06.002 (2004).

10 Bernhardt, T. G. & de Boer, P. A. J. SlmA, a Nucleoid-Associated, FtsZ Binding Protein Required for Blocking Septal Ring Assembly over Chromosomes in E. coli. Molecular Cell 18, 555–564, doi:10.1016/j.molcel.2005.04.012 (2005).

11 de Boer, P. A., Crossley, R. E. & Rothfield, L. I. A division inhibitor and a topological specificity factor coded for by the minicell locus determine proper placement of the division septum in E. coli. Cell 56, 641–649, doi:10.1016/0092-8674(89)90586-2 (1989).

12 Pang, T., Wang, X., Lim, H. C., Bernhardt, T. G. & Rudner, D. Z. The nucleoid occlusion factor Noc controls DNA replication initiation in Staphylococcus aureus. PLOS Genetics 13, e1006908, doi:10.1371/journal.pgen.1006908 (2017).

13 Veiga, H., Jorge, A. M. & Pinho, M. G. Absence of nucleoid occlusion effector Noc impairs formation of orthogonal FtsZ rings during Staphylococcus aureus cell division. Molecular Microbiology 80, 1366–1380, doi:https://doi.org/10.1111/j.1365-2958.2011.07651.x (2011).

14 Tzagoloff, H. & Novick, R. Geometry of Cell Division in Staphylococcus aureus. Journal of Bacteriology 129, 343–350 (1976).

15 Turner, R. D. et al. Peptidoglycan architecture can specify division planes in Staphylococcus aureus. Nat Commun 1, 26, doi:10.1038/ncomms1025 (2010).

16 Saraiva, B. M. et al. Reassessment of the distinctive geometry of Staphylococcus aureus cell division. Nature Communications 11, doi:10.1038/s41467-020-17940-9 (2020).

17 Berney, M., Hammes, F., Bosshard, F., Weilenmann, H. U. & Egli, T. Assessment and interpretation of bacterial viability by using the LIVE/DEAD BacLight Kit in combination with flow cytometry. Appl Environ Microbiol 73, 3283–3290, doi:10.1128/aem.02750-06 (2007).

18 Bouvier, T., Troussellier, M., Anzil, A., Courties, C. & Servais, P. Using light scatter signal to estimate bacterial biovolume by flow cytometry. Cytometry 44, 188–194, doi:10.1002/1097-0320(20010701)44:3<188::aid-cyto1111>3.0.co;2-c (2001).

19 Laubacher, M. E., Melquist, A. L., Chandramohan, L. & Young, K. D. Cell sorting enriches Escherichia coli mutants that rely on peptidoglycan endopeptidases to suppress highly aberrant morphologies. J Bacteriol 195, 855–866, doi:10.1128/jb.01450-12 (2013).

20 Nega, M., Tribelli, P. M., Hipp, K., Stahl, M. & Götz, F. New insights in the coordinated amidase and glucosaminidase activity of the major autolysin (Atl) in Staphylococcus aureus. Communications Biology 3, 695, doi:10.1038/s42003-020-01405-2 (2020).

21 Wheeler, R. et al. Bacterial Cell Enlargement Requires Control of Cell Wall Stiffness Mediated by Peptidoglycan Hydrolases. mBio 6, e00660, doi:10.1128/mBio.00660-15 (2015).

22 Chan, Y. G., Frankel, M. B., Missiakas, D. & Schneewind, O. SagB Glucosaminidase Is a Determinant of Staphylococcus aureus Glycan Chain Length, Antibiotic Susceptibility, and Protein Secretion. J Bacteriol 198, 1123–1136, doi:10.1128/jb.00983-15 (2016).

23 Schaefer, K. et al. Structure and reconstitution of a hydrolase complex that may release peptidoglycan from the membrane after polymerization. Nature Microbiology 6, 34–43, doi:10.1038/s41564-020-00808-5 (2021).

24 Hsu, Y. P. et al. Full color palette of fluorescent d-amino acids for in situ labeling of bacterial cell walls. Chem Sci 8, 6313–6321, doi:10.1039/c7sc01800b (2017).

25 Meeske, A. J. et al. High-Throughput Genetic Screens Identify a Large and Diverse Collection of New Sporulation Genes in Bacillus subtilis. PLOS Biology 14, e1002341, doi:10.1371/journal.pbio.1002341 (2016).

26 Zhou, X. et al. Mechanical crack propagation drives millisecond daughter cell separation in Staphylococcus aureus. Science 348, 574–578, doi:10.1126/science.aaa1511 (2015).

27 Jorge, A. M., Hoiczyk, E., Gomes, J. P. & Pinho, M. G. EzrA Contributes to the Regulation of Cell Size in Staphylococcus aureus. PLOS ONE 6, e27542, doi:10.1371/journal.pone.0027542 (2011).

28 Steele, V. R., Bottomley, A. L., Garcia-Lara, J., Kasturiarachchi, J. & Foster, S. J. Multiple essential roles for EzrA in cell division of Staphylococcus aureus. Molecular Microbiology 80, 542–555, doi:10.1111/j.1365-2958.2011.07591.x (2011).

29 Levin, P. A., Kurtser, I. G. & Grossman, A. D. Identification and characterization of a negative regulator of FtsZ ring formation in Bacillus subtilis. Proceedings of the National Academy of Sciences 96, 9642–9647, doi:10.1073/pnas.96.17.9642 (1999).

30 Andreu, J. M. et al. The antibacterial cell division inhibitor PC190723 is an FtsZ polymer-stabilizing agent that induces filament assembly and condensation. J Biol Chem 285, 14239–14246, doi:10.1074/jbc.M109.094722 (2010).

31 Halbedel, S. & Lewis, R. J. Structural basis for interaction of DivIVA/GpsB proteins with their ligands. Molecular Microbiology 111, 1404–1415, doi:https://doi.org/10.1111/mmi.14244 (2019).

32 Eswara, P. J. et al. An essential Staphylococcus aureus cell division protein directly regulates FtsZ dynamics. eLife 7, doi:10.7554/elife.38856 (2018).

33 Hammond, L. R. et al. GpsB Coordinates Cell Division and Cell Surface Decoration by Wall Teichoic Acids in Staphylococcus aureus. Microbiology Spectrum, doi:10.1128/spectrum.01413-22 (2022).

34 Coe, K. A. et al. Multi-strain Tn-Seq reveals common daptomycin resistance determinants in Staphylococcus aureus. PLOS Pathogens 15, e1007862, doi:10.1371/journal.ppat.1007862 (2019).

35 Fey, P. D. et al. A Genetic Resource for Rapid and Comprehensive Phenotype Screening of Nonessential Staphylococcus aureus Genes. mBio 4, e00537–00512, doi:10.1128/mBio.00537-12 (2013).

36. Sacco, M. D., et al. Staphylococcus aureus FtsZ and PBP4 bind to the conformationally dynamic N-terminal domain of GpsB. bioRxiv, doi:10.1101/2022.10.25.513704 (2022).

37 Datsenko, K. A. & Wanner, B. L. One-step inactivation of chromosomal genes in Escherichia coli K-12 using PCR products. Proceedings of the National Academy of Sciences 97, 6640–6645, doi:10.1073/pnas.120163297 (2000).

38 Liew, A. T. F. et al. A simple plasmid-based system that allows rapid generation of tightly controlled gene expression in Staphylococcus aureus. Microbiology 157, 666–676, doi:https://doi.org/10.1099/mic.0.045146-0 (2011).

39 Deatherage, D. E. & Barrick, J. E. Identification of mutations in laboratory-evolved microbes from next-generation sequencing data using breseq. Methods Mol Biol 1151, 165–188, doi:10.1007/978-1-4939-0554-6_12 (2014).

40 Ducret, A., Quardokus, E. M. & Brun, Y. V. MicrobeJ, a tool for high throughput bacterial cell detection and quantitative analysis. Nature Microbiology 1, 16077, doi:10.1038/nmicrobiol.2016.77 (2016).

41 Dobihal, G. S., Brunet, Y. R., Flores-Kim, J. & Rudner, D. Z. Homeostatic control of cell wall hydrolysis by the WalRK two-component signaling pathway in Bacillus subtilis. eLife 8, e52088, doi:10.7554/eLife.52088 (2019).

42 Mendler, K. et al. AnnoTree: visualization and exploration of a functionally annotated microbial tree of life. Nucleic Acids Research 47, 4442–4448, doi:10.1093/nar/gkz246 (2019).

43 Sievers, F. et al. Fast, scalable generation of high-quality protein multiple sequence alignments using Clustal Omega. Molecular Systems Biology 7, 539, doi:https://doi.org/10.1038/msb.2011.75 (2011).

44 Omasits, U., Ahrens, C. H., Müller, S. & Wollscheid, B. Protter: interactive protein feature visualization and integration with experimental proteomic data. Bioinformatics 30, 884–886, doi:10.1093/bioinformatics/btt607 (2014).

45 Lupas, A., Van Dyke, M. & Stock, J. Predicting coiled coils from protein sequences. Science 252, 1162–1164, doi:10.1126/science.252.5009.1162 (1991).

46 Erdős, G. & Dosztányi, Z. Analyzing Protein Disorder with IUPred2A. Current Protocols in Bioinformatics 70, e99, doi:https://doi.org/10.1002/cpbi.99 (2020).

47 Jumper, J. et al. Highly accurate protein structure prediction with AlphaFold. Nature 596, 583–589, doi:10.1038/s41586-021-03819-2 (2021).

