## Supplemental Information for "Identification of FacZ as a division site placement factor in *Staphylococcus aureus*"

**Affiliations:**
[1]Department of Microbiology, Blavatnik Institute, Harvard Medical School, Boston, Massachusetts, USA
[2]Department of Biochemistry & Biophysics, School of Medicine, University of North Carolina at Chapel Hill, Chapel Hill, NC, USA
[3]Lineberger Comprehensive Cancer Center, University of North Carolina at Chapel Hill, Chapel Hill, NC, USA
[4]Howard Hughes Medical Institute, Harvard Medical School, Boston, Massachusetts, USA

***To whom correspondence should be addressed:**

Thomas G. Bernhardt, Ph.D.
Harvard Medical School
Department of Microbiology
Boston, Massachusetts 02115


David Z. Rudner
Harvard Medical School
Department of Microbiology
Boston, Massachusetts 02115, USA


### Supplemental Figures and Legends

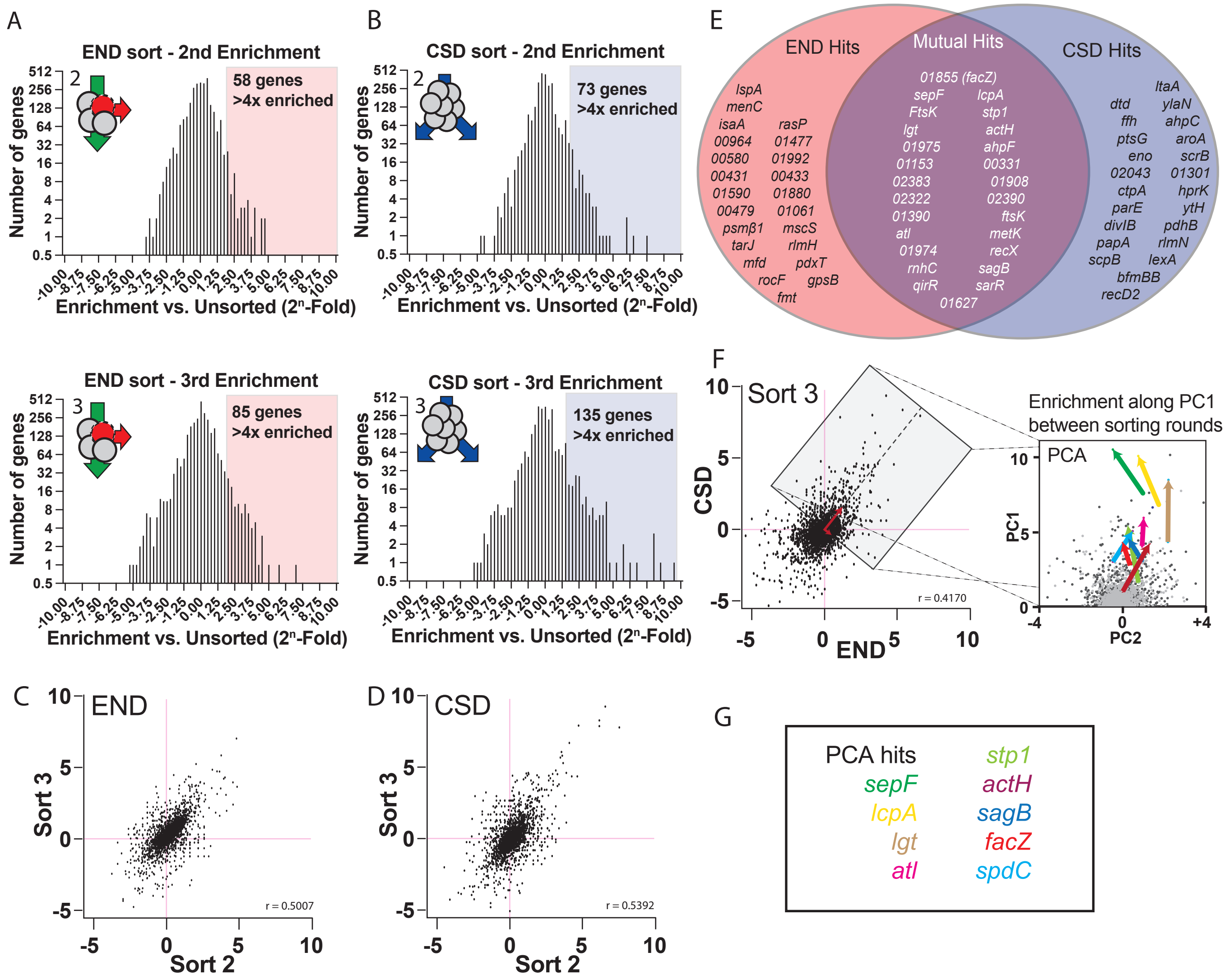


**Figure S1. CSD and END enrichment data.**

(A-B) Histograms showing the relative enrichment of transposon insertions in genes following the second and third END (A) or CSD (B) sorting rounds. Colored boxes highlight genes with >4x enrichment in transposon insertions. (C-D) Scatterplots showing relative enrichment at each locus in sort 3 over relative enrichment at the same locus in sort 2 for both the END (C) and CSD

(D) screens. Linear regression analysis indicates that relative enrichment is well-correlated between sorting rounds, suggesting enrichment is driven by phenotypic selection and not chance. (E) Venn diagram comparison of the 50 genes in which transposon insertions were most strongly-enriched in the CSD and END sorts. (F) Enrichment in the final CSD sort is moderately correlated with enrichment in the final END sort, consistent with the overlapping mutant populations isolated from the two screens. 2D-PCA of the final sorts identified a vector (PC1, red) that serves as a proxy for enrichment in both sorts, allowing ranking of mutual hits based on enrichment along this vector between rounds of sorting (PCA panel, and see **Table S4-5**). The PCA pop-out panel (right) shows relative enrichment at each genetic locus in round 2 (light gray) and round 3 (dark gray), rotated so that PC1 is vertical. Colored arrows show the change in enrichment between rounds of sorting for a subset of hits with known roles in envelope biogenesis. Upward movement in this vector space indicates enrichment in both screens. (G) A selection of hits identified in the PCA, many of which are known cell-envelope biogenesis genes. Text colors match with the arrow colors for the given genes in the PCA plot.

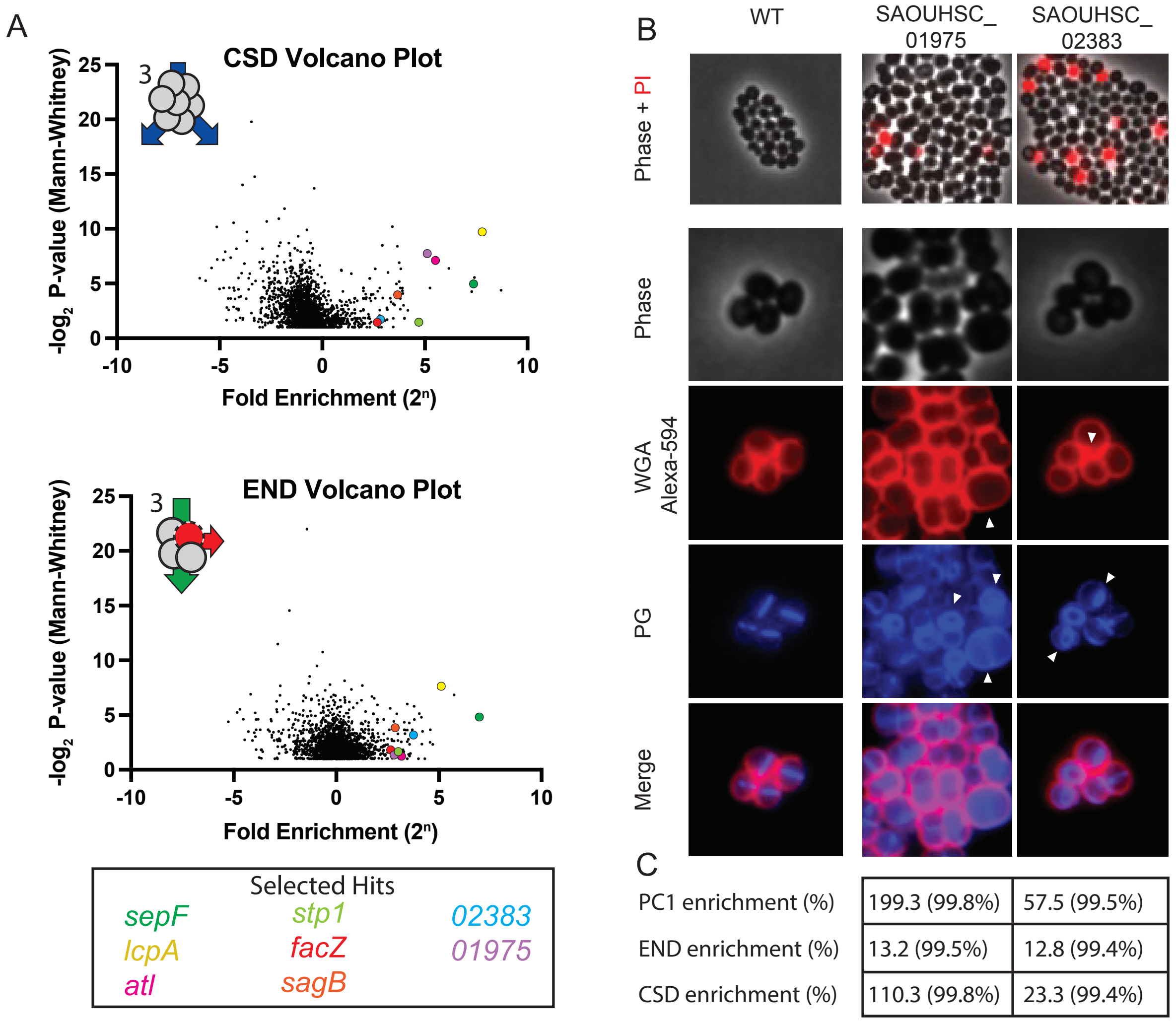


| | SAOUHSC_01975 | SAOUHSC_02383 |
|---|---|---|
| PC1 enrichment (%) | 199.3 (99.8%) | 57.5 (99.5%) |
| END enrichment (%) | 13.2 (99.5%) | 12.8 (99.4%) |
| CSD enrichment (%) | 110.3 (99.8%) | 23.3 (99.4%) |

**Figure S2. Validation and characterization of additional hits from the screen.**

A) Volcano plots showing the relative enrichment of transposon insertions in each locus versus the p-value from Mann-Whitney U for the same locus. The final CSD sort (top) and final END sort (middle) are compared to an ungated control generated in parallel. Selected hits (bottom) are color coded and correspond to the colored circle in the plots. B) Representative images of two mutants identified in our screens, derived from the NTML Tn-mutant library. To assess END phenotype (top), mid-log phase cells were washed with PBS and incubated for 5 minutes with PI, then placed on pads containing PI and imaged. To assess CSD phenotype (bottom), mid-log phase cells were pulse-labeled with HADA to visualize peptidoglycan synthesis (PG) (middle row, blue), washed three times with PBS to arrest growth and remove incorporated HADA, and then labeled for with WGA Alexa-594 to label teichoic acid (top row, red). Compared to the parental *S. aureus* USA300 strain (WT, aTB001), Tn-inactivation of USA300 orthologs of SAOUHSC_01974 (aTB112) and SAOUHSC_02383 (aTB113), have increased PI staining and morphological defects (white carets), consistent with END and CSD phenotypes. C) Table showing the fold-change of transposon insertions in the genes inactivated in (B) in the final round of the END and CSD sorts, as well as relative enrichment along PC1. The enrichment percentile of mutations in those genes compared to all mutants is given in parentheses.

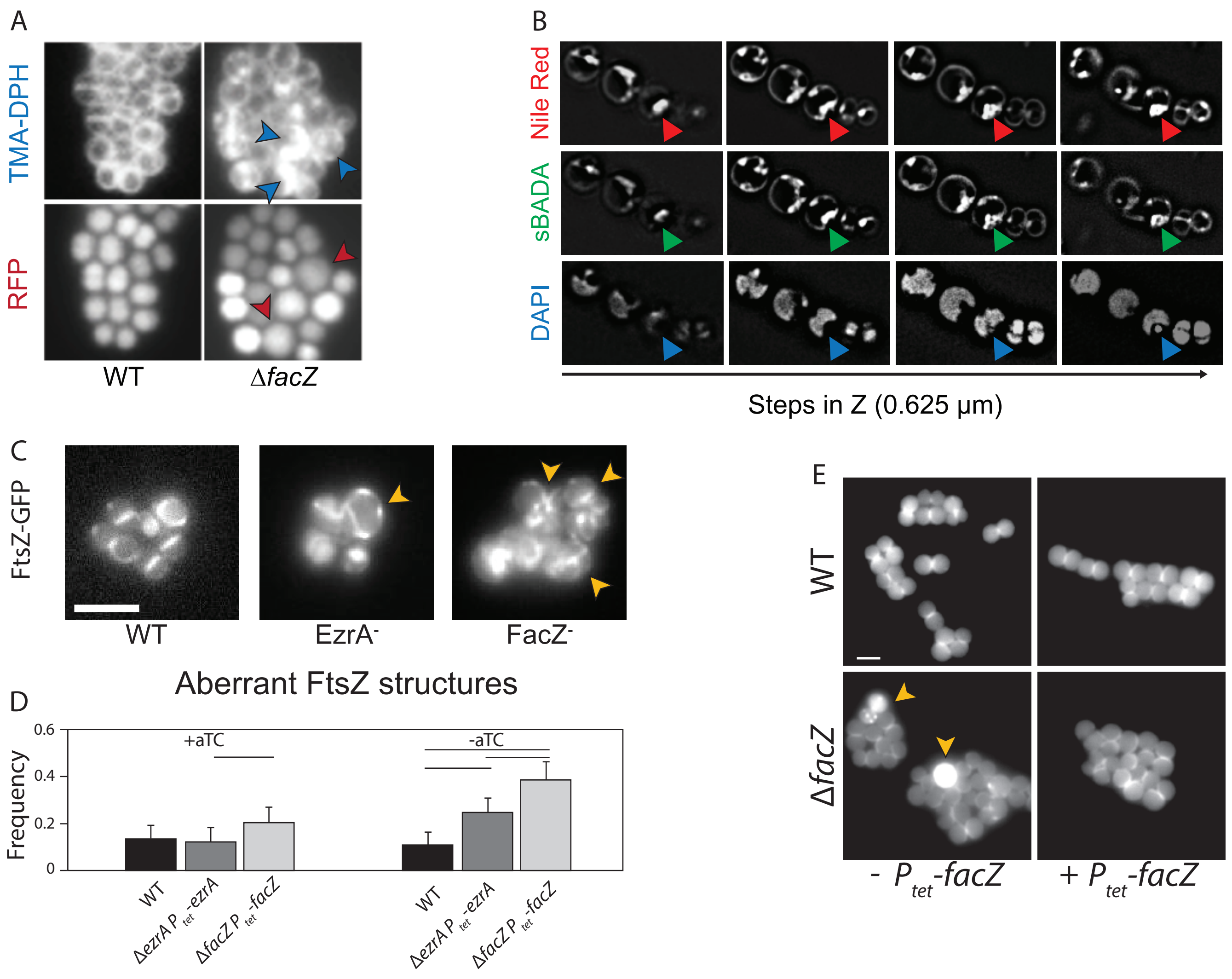


**Figure S3. Inactivation of FacZ impairs cell division and morphogenesis.**

A) Representative images showing membrane labeling and cytoplasmic fluorescence of WT (aTB523) and Δ*facZ* (aTB527) cells used to quantify morphological and membrane defects in **Fig. 3B-C**. Cells were grown overnight in the presence of TMP to maintain the RFP-bearing plasmid, then subcultured into medium free of antibiotics, and grown into mid-log phase. These cells were then labeled with TMA-DPH, imaged on M9 pads (2% agarose), and segmented on cytoplasmic fluorescence signal; cell size measurements for violin plots (**Fig. 3B**) were automated using MicrobeJ; red carets point to unusually large cells. For each cell identified by MicrobeJ, the number of aberrant membrane foci (blue carets) was recorded manually (**Fig. 3C**). B) Z-stack of 3D-SIM reconstruction of *ΔfacZ* cells (aTB251) stained identically to **Fig. 3** shows aberrant features. Red carets follow membrane features (Nile Red, top) through Z-planes (left to right); features are continuous throughout the cell in the Z-dimension, and do not emerge in the middle of the cytoplasm, consistent with these membrane features being continuous invaginations of the cell membrane, rather than completely internalized structures. Analogous continuous features are apparent when imaging labeled cell wall (green carets, sBADA, middle), and correspond to local exclusion of the nucleoid (blue carets, DAPI, bottom), both of which also extend throughout all Z-slices. C) FtsZ-GFP was induced at low levels in exponentially-growing *S. aureus* to determine which cells were dividing and to determine the orientation of the division plane. In WT *S. aureus* (aTB219), most cells exhibited a single-ring, consistent with normal cell division. In cells depleted of FacZ (aTB390, FacZ$^{-}$), aberrant FtsZ structures (defined as multiple Z-structures, drastically off-center Z-structures, or diffuse cytoplasmic FtsZ-GFP signal) were apparent. A similar phenotype was observed in cells depleted for the division protein EzrA (aTB391, EzrA$^{-}$). D) Bar graphs showing quantification of aberrant FtsZ structures ($n \geq 100$ cells per condition), with error bars displaying 95% confidence intervals and horizontal bars indicating p-value <0.001 (chi-squared test). The defect was largely rescued in the presence of aTC (25 ng/mL). E) WT *S. aureus* with (aTB341) and without (aTB003) an integrated *pTP63-facZ* expression construct were imaged alongside Δ*facZ* cells with (aTB372) and without (aTB251) the same integrated *facZ* expression construct. Cells were grown to mid-log phase, treated with aTC (25 ng/mL) to induce FacZ expression from the integrated $P_{tet}$ promoter, and exposed to HADA to label active zones of PG insertion. Only Δ*facZ* cells lacking the complementing allele exhibited morphological defects (yellow carets) (scale bar = 2 µm).

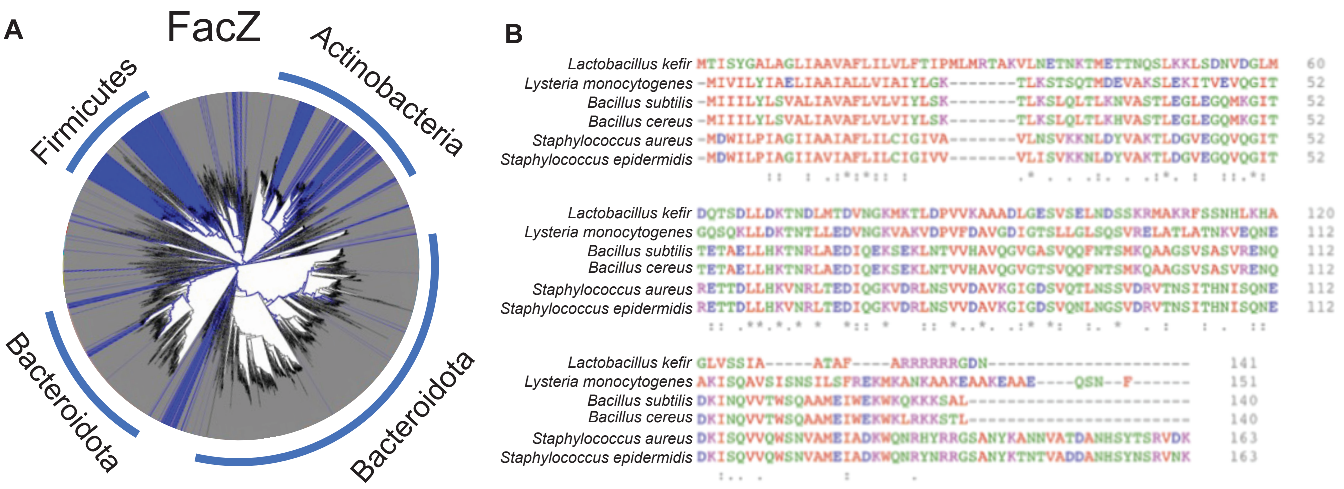


**Figure S4. Conservation of FacZ.**

A) Dendrogram highlighting the distribution of FacZ in a broad range of bacterial species. B) Alignment of FacZ orthologs from selected Firmicutes.

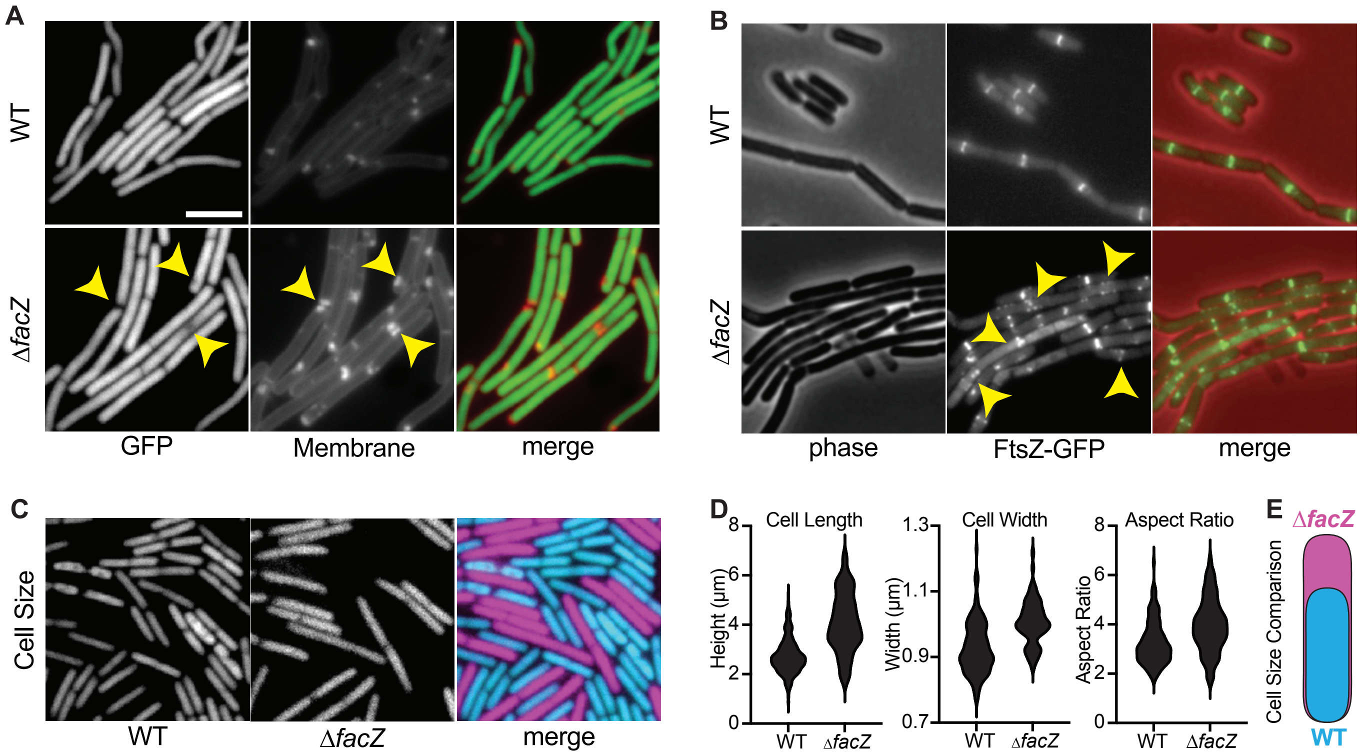


**Figure S5. FacZ is required for envelope biogenesis and cell division in *B. subtilis.***

All microscopy was performed on exponentially-growing derivatives of *B. subtilis* strain PY79 imaged on agarose pads. All images were scaled identically (scale bar = 4 µm). A) WT (bDR2789) and Δ*facZ* (bTB039) cells that constitutively express cytoplasmic GFP were labeled with the membrane dye Nile Red. Yellow carets highlight membrane invaginations (membrane, middle) that correspond to local depletions of cytoplasmic GFP (left). These phenotypes are similar to those associated with Δ*facZ* in *S. aureus* (**Fig. 3**). B) WT (bDR2229) and Δ*facZ* (bTB018) cells expressing FtsZ-GFP from an ectopic locus.  Aberrant FtsZ structures are highlighted with yellow carets. C) WT (bDR2637, false-colored cyan) cells expressing RFP and Δ*facZ* (bTB040, false-colored magenta) cells expressing BFP were co-cultured and imaged on the same pads, allowing for direct comparison of cell size. D) Cells were segmented using cytoplasmic fluorescence in MicrobeJ to compare cell morphology. Violin plots show cell length, cell width, and aspect ratio of WT and Δ*facZ* cells from a representative experiment (n >100 cells). Dotted lines show median and solid lines show quartiles. E) Cartoon depicting the morphological differences between a typical WT and Δ*facZ* cell using the median parameters determined in this analysis.

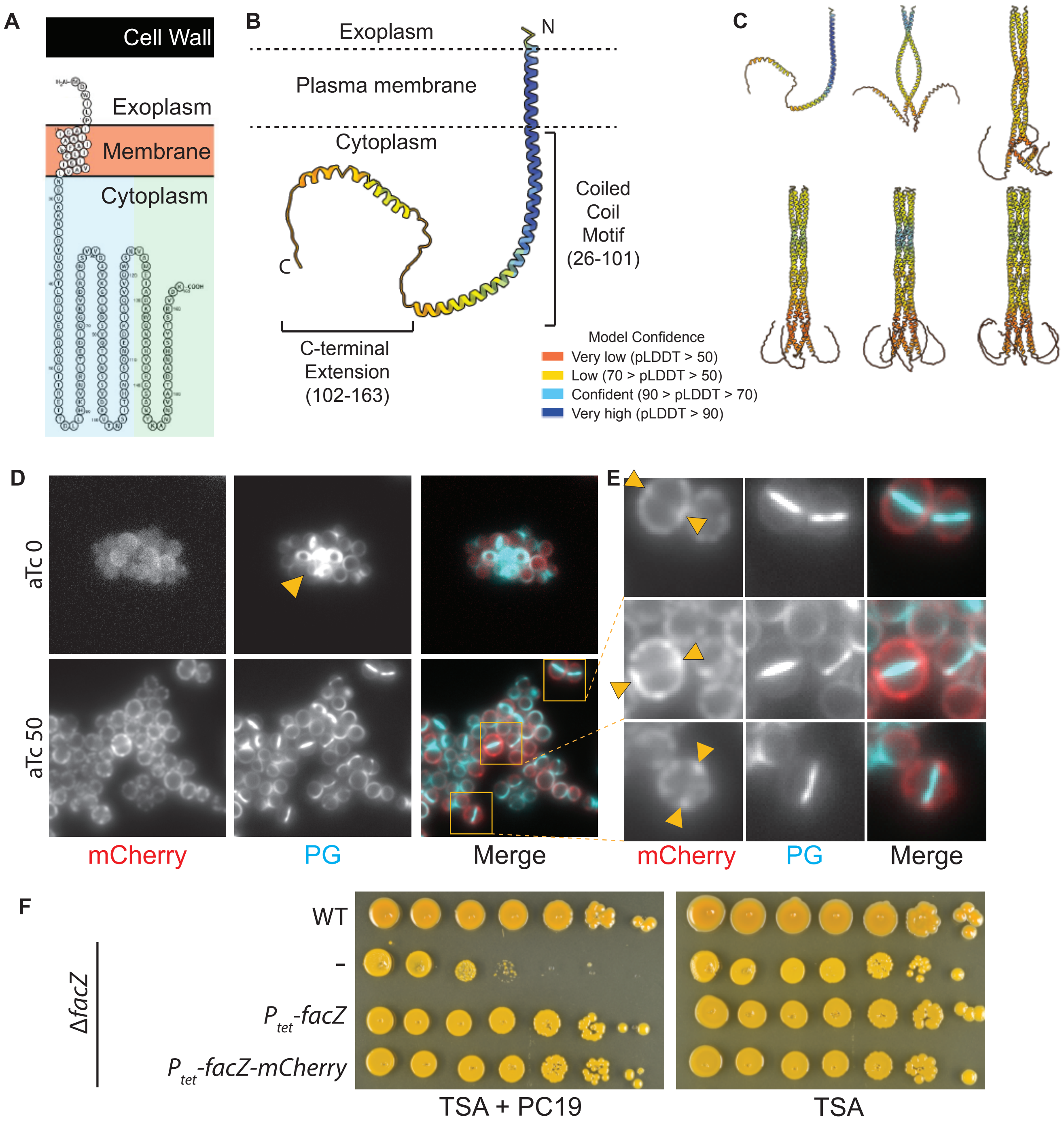


**Figure S6. FacZ structural predictions and functionality of the FacZ-mCherry fusion.**

A) Predicted membrane topology of FacZ (Protter)[1]. FacZ is predicted to have one transmembrane helix with a short extracellular N-terminal region. Most of the protein is predicted to extend into the cytoplasm, with a coiled-coil region (blue box) and a disordered C-terminal extension (green box). B) AlphaFold2 model of full-length FacZ, colored by pLDDT confidence score. Predicted domains and orientation in the plasma membrane are labeled. C) AlphaFold2-multimer predictions of FacZ assemblies from one copy to six copies. D) A Δ*facZ* strain harboring *facZ-mCherry* under control of a $P_{tet}$ promoter (aTB373) was grown into mid-log phase with or without aTC and pulse-labeled with HADA and imaged on PBS pads containing 2% agarose. Cells depleted of FacZ-mCherry (red) had aberrant sites of PG synthesis (yellow caret) consistent with inactivation of FacZ, while aTC (50 ng/mL) induction of *facZ-mCherry* restored normal morphology and PG incorporation. E) When FacZ-mCherry is expressed at levels that restore normal cell morphology to Δ*facZ*, the fluorescent fusion is enriched at periseptal regions, flanking actively-growing zones of septal PG synthesis (yellow carets). F) WT (aTB003), Δ*facZ* (aTB251), and Δ*facZ* cells expressing untagged *facZ* (aTB372) or *facZ-mCherry* (aTB373) under the control of the $P_{tet}$ promoter were normalized to $OD_{600}$ = 1.0, serially diluted, and spotted onto TSA agar plates with aTC (50ng/mL) and with or without PC190723 (100 ng/mL) as indicated.

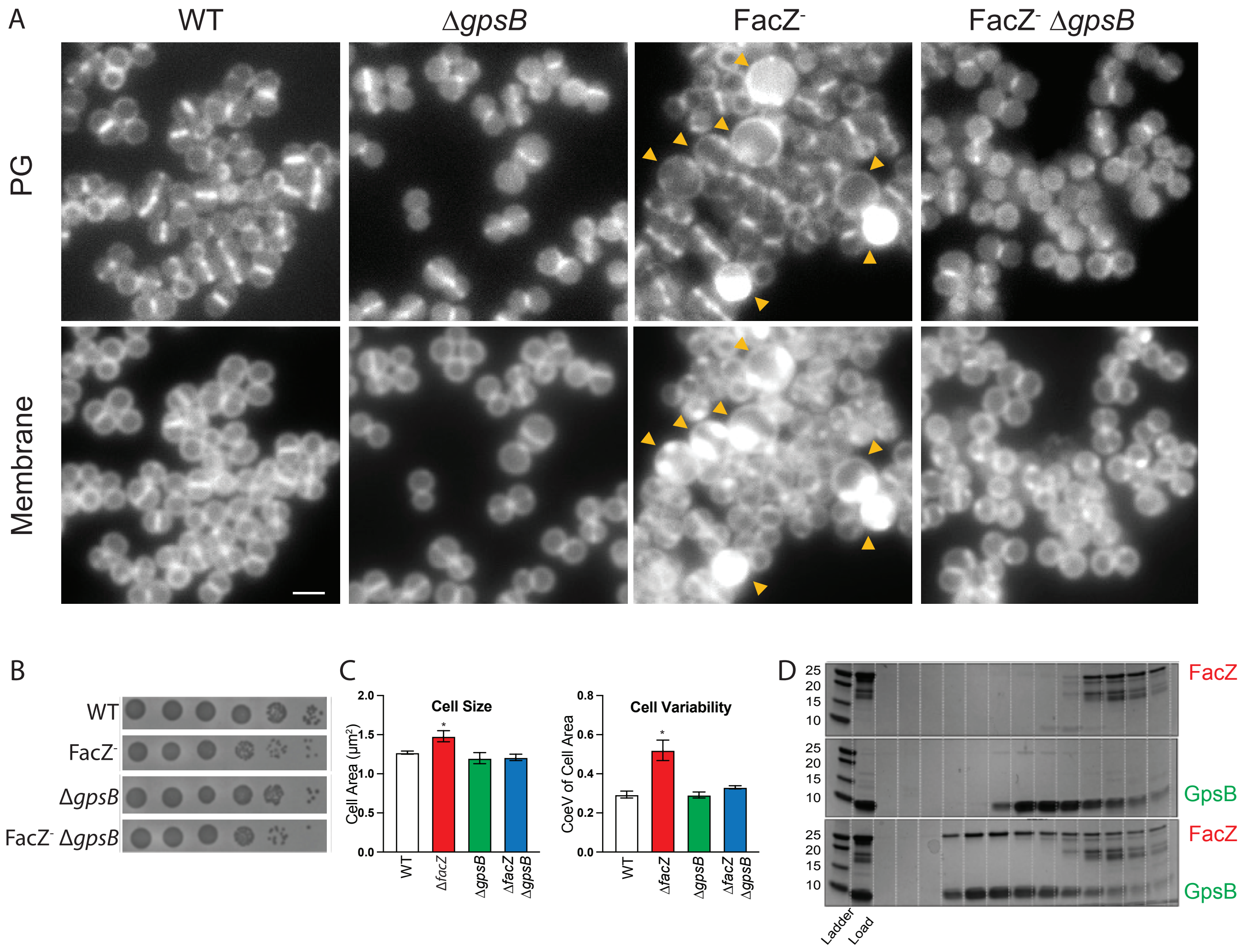


**Figure S7. Δ*gpsB* corrects the morphological defects of Δ*facZ* mutants.**

A) Representative images of WT (aTB003), Δ*gpsB* (aTB525), FacZ- (aTB372, *ΔfacZ pTP63d-facZ*), and FacZ- Δ*gpsB* (aTB540, *ΔfacZ pTP63d-facZ ΔgpsB*) grown into mid-log phase without induction of *facZ*, and pulse-labeled with sBADA to label active PG insertion and Nile Red to label cell membranes (see **Fig. 5E**). Inactivation of *gpsB* has minimal impact on cell morphology or localization of envelope probes. Depletion of FacZ causes characteristic envelope and morphological defects (yellow carets), which are rescued by inactivation of *gpsB*. B) Spot titers of the same strains imaged in panel A confirms that depletion of FacZ and/or inactivation of *gpsB* has negligible impact on cell viability. C) Median cell size and coefficient of variance of cell size (CoeV) of the indicated strains from replicated experiments (see pooled data in **Fig. 5F**).

Inactivation of *gpsB* alone has no significant impact on size or variability, but rescues defects caused by depletion of FacZ (Student's t-test, p < 0.01). D) Representative SDS-PAGE gels of fractions from the size-exclusion column from runs with FacZ alone (top), GpsB alone (middle), and a mixture of both proteins (bottom).

**Supplemental Video 1***

Video advancing through Z-stack of 3D-SIM reconstruction of Δ*facZ* cells (see **Fig. S3B**).

### Supplemental Tables and Legends

| **Table S1: Tn-seq meta-analysis** | | All Unique TAs | | Unique Mapped TAs | | | Genes Hit | | | | | | | | |
|---|---|---|---|---|---|---|---|---|---|---|---|---|---|---|---|
| | | | | Count | Per gene | | | | | | | | | | |
| Symbol | Sort | Count | % | | Median | Mean | 1+ | % | vs NTML | 2+ | % | vs NTML | 5+ | % | vs NTML |
| 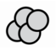 | Initial | 60485 | 22.4 | 56134 | 17 | 21.4 | 2468 | 93.9 | 1.258 | 2330 | 88.6 | 1.188 | 2045 | 77.8 | 1.042 |
| | Ungated 2 | 45082 | 16.7 | 39910 | 12 | 15.2 | 2438 | 92.7 | 1.243 | 2272 | 86.4 | 1.158 | 1861 | 70.8 | 0.949 |
| | Ungated 3 | 40682 | 15.1 | 35566 | 10 | 13.5 | 2415 | 91.9 | 1.231 | 2230 | 84.8 | 1.137 | 1791 | 68.1 | 0.913 |
| 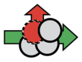 | END 2 | 36553 | 13.6 | 31664 | 9 | 12 | 2388 | 90.8 | 1.217 | 2193 | 83.4 | 1.118 | 1750 | 66.6 | 0.892 |
| | END 3 | 34832 | 12.9 | 29590 | 8 | 11.3 | 2429 | 92.4 | 1.238 | 2215 | 84.3 | 1.129 | 1739 | 66.1 | 0.886 |
| 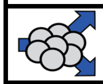 | CSD 2 | 40631 | 15.1 | 35729 | 10 | 13.6 | 2408 | 91.6 | 1.227 | 2201 | 83.7 | 1.122 | 1789 | 68 | 0.912 |
| | CSD 3 | 32378 | 12 | 27554 | 8 | 10.5 | 2367 | 90 | 1.206 | 2140 | 81.4 | 1.091 | 1652 | 62.8 | 0.842 |

**Table S1. Tn-seq meta-analysis.**

Table summarizes the relevant metrics from Tn-Seq analysis of the initial transposon library, END and CSD enrichments, and the ungated control sort. The numbers 2 and 3 refer to the sorting round (e.g., CSD 2 has been subjected to two rounds of sorting increased light scattering). The total number of unique TA sites and the percentage of total TA sites in the genome with transposon insertions are given, as well as median and mean number of insertions per gene in the library following enrichments. The number of genes in each library hit at least once, twice, or five times is shown at right, along with the percent of annotated genes bearing at least that many hits. Also, the number of genes meeting or exceeding these thresholds is compared to the number of mutants in the NTML ordered transposon library[2], which is commonly used in *S. aureus* research.

**Table S2. Tn-seq data for the END and CSD enrichments***

Tn-seq results for the second and third rounds of sorting of the CSD and END enrichments. Relative enrichments compare the results of a given round of either CSD or END sorting to the ungated control. P-values are from Mann-Whitney U test. Genes are sorted by SAOUHSC locus number.

**Table S3. Relative enrichment of transposon insertions in each gene following END and CSD enrichments versus the ungated control***

Tn-seq data showing the relative enrichments of the END and CSD sorts compared to an ungated control. Genes are sorted by SAOUHSC locus number.

**Table S4. PCA-adjusted Tn-Seq results***

Following 2D-PCA rotation of the CSD and END sorted data, each gene's enrichment was assigned a new (X,Y) location, with X representing a genes relative enrichment along PC2, and Y representing the relative enrichment along PC1. These data were used to plot the PCA panel in **Fig. S1**. Genes are sorted by SAOUHSC locus number.

**Table S5. PCA-adjusted hit list.**

The top 50 hits from the PCA analysis based on maximum relative enrichment along PC1. "Δ" indicates the change in PC1 or PC2 relative enrichment between rounds of sorting. "Mean fold increase" and "% increasing in enrichment" are metrics to assess whether hits identified by PCA in sort 2 increase in PC1 enrichment from additional rounds of CSD and END sorting. In general, hits identified by PC1 enrichment enrich further from additional sorts, where those from PC2 do not, consistent with PC1-enrichment serving as a proxy for mutual CSD and END enrichment. Genes are displayed by decreasing relative enrichment along PC1.

**Table S6. Δ*facZ* synthetic lethal Tn-seq***

Tn-seq analysis of transposon libraries constructed in WT *S. aureus* (aTB015) versus one constructed in a Δ*facZ* (aTB259) strain. P-values are from Mann-Whitney U test. Genes are sorted by SAOUHSC locus number. Genes are sorted by increasing Δ*facZ:WT* relative enrichment ratio.

* - Due to formatting restrictions, videos and large data sets will be made available upon publication**.**

**Table S7: *S. aureus* strains used in this study**

| ***S. aureus* strain** | **Background** | **Relevant genotype** | **Source** | **Construction notes** |
|---|---|---|---|---|
| **aTB001** | USA300 | WT | Pang, T. et al.[3] | - |
| **aTB003** | HG003 | WT | Pang, T. et al.[3] | - |
| **aTB004** | RN4220 | WT | Pang, T. et al.[3] | - |
| **aTB015** | RN4220 | Δ*attB(f11)::Orf5, pTM378* | Wang, H. et al.[4] | - |
| **aTB016** | RN4220 | Δ*attB(f11)::Orf5, pTM381* | Wang, H. et al.[4] | - |
| **aTB033** | RN4220 | pTP044 | Pang, T. et al.[3] | - |
| **aTB111** | USA300 | *Tn5::USA300_0932 (SAOUHSC_00965)* | Fey, P. et al.[2] | - |
| **aTB112** | USA300 | *Tn5::USA300_1792 (SAOUHSC_01975)* | Fey, P. et al.[2] | - |
| **aTB113** | USA300 | *Tn5::USA300_2094 (SAOUHSC_02383)* | Fey, P. et al.[2] | - |
| **aTB219** | HG004 | *pLOW-ftsZ-GFP* | This study | pLOW-ftsZ-GFP transduced into aTB003 |
| **aTB243** | RN4220 | Δ*facZ::specR* | This study | Homologous recombination of pMR91-Δ*facZ::specR* into aTB004 |
| **aTB251** | HG003 | Δ*facZ::specR* | This study | Φ85 lysate of aTB243 was used to transduce Δ*facZ::specR* into aTB003 |
| **aTB259** | RN4220 | Δ*facZ::specR* Δ*attB(f11)::Orf5, pTM378* | This study | Φ85 lysate of aTB243 was used to transduce Δ*facZ::specR* into aTB015 |
| **aTB261** | RN4220 | Δ*facZ::specR* Δ*attB(f11)::Orf5, pTM381* | This study | Φ85 lysate of aTB243 was used to transduce Δ*facZ::specR* into aTB016 |
| **aTB263** | RN4220 | *pGL485* | This study | pGL485 was transformed into aTB004 |
| **aTB287** | HG003 | Δ*sagB (lytD)* | This study | Provided by Walker lab |
| **aTB315** | RN4220 | *pGL485 pLOW-facZ* | This study | *pLOW-facZ* was transformed into aTB263 |
| **aTB316** | RN4220 | *pGL485 pLOW-facZ-mCherry* | This study | *pLOW-facZ-mCherry* was transformed into aTB263 |
| **aTB339** | RN4220 | *pTP044 pTP63d-facZ* | This study | *pTP63d-facZ* was transformed into aTB044 |
| **aTB341** | HG003 | *pTP63d-facZ* | This study | Φ85 lysate of aTB339 was used to transduce *pTP63-facZ* into aTB003 |
| **aTB347** | HG003 | *pLOW-facZ* | This study | Φ85 lysate of aTB315 was used to transduce *pLOW-facZ* into aTB003 |
| **aTB348** | HG003 | *pLOW-facZ-mCherry* | This study | Φ85 lysate of aTB339 was used to transduce *pTP63-facZ-mCherry* into aTB003 |
| **aTB356** | HG003 | Δ*facZ::specR pLOW-facZ* | This study | Φ85 lysate of aTB243 was used to transduce Δ*facZ::specR* into aTB347 |
| **aTB358** | HG003 | Δ*facZ::specR pLOW-facZ-mCherry* | This study | Φ85 lysate of aTB243 was used to transduce Δ*facZ::specR* into aTB348 |
| **aTB372** | HG003 | Δ*facZ::specR pTP63d-facZ* | This study | Φ85 lysate of aTB243 was used to transduce Δ*facZ::specR* into aTB341 |
| **aTB374** | RN4220 | *pTP044 pTP63d-facZ-mCherry* | This study | *pTP63d-facZ-mCherry* was transformed into aTB033 |
| **aTB376** | RN4220 | *pTP044 pTP63-ftsZ-gfp* | This study | *pTP63-ftsZ-gfp* was transformed into aTB033 |
| **aTB378** | HG003 | Δ*ezrA pTP63b-ezrA* Δ*facZ::specR* | This study | Φ85 lysate of aTB243 was used to transduce Δ*facZ::specR* into aTB481 |
| **aTB390** | HG003 | Δ*facZ::specR pTP63d-facZ pLOW-FtsZ-GFP* | This study | Φ85 lysate of *RN4220 pLOW-ftsZ-GFP* was transduced into aTP372 |
| **aTB391** | HG003 | Δ*ezrA pTP63d-ezrA pLOW-FtsZ-GFP* | This study | Φ85 lysate of *RN4220 pLOW-ftsZ-GFP* was transduced into aTP481 |
| **aTB392** | HG003 | *pTP63d-facZ-mCherry* | This study | Φ85 lysate of aTB374 used to transduce *pTP63-facZ-mCherry* into aTB003 |
| **aTB394** | HG003 | *pTP63d-ftsZ-GFP* | This study | Φ85 lysate of aTB376 used to transduce *pTP63-ftsZ-GFP* into aTB003 |

| | | | | |
|---|---|---|---|---|
| **aTB453** | HG003 | *ΔfacZ::specR gpsB (T)6→5 (truncation 187)* | This study | Spontaneous suppressor of aTB251 sensitivity to PC190723 |
| **aTB476** | HG003 | *ΔfacZ::specR gpsB(Y26*)* | This study | Spontaneous suppressor of aTB251 sensitivity to PC190723 |
| **aTB478** | HG003 | *ΔfacZ::specR gpsB (T)6-5 (trunc. 128)* | This study | Spontaneous suppressor of aTB251 sensitivity to PC190723 |
| **aTB492** | HG003 | *gpsB::Tn5* | This study | Φ85 lysate of NTML strain *gpsB::Tn5(erm)* transduced into aTB003 |
| **aTB497** | HG003 | *gpsB::Tn5 ΔfacZ::specR* | This study | Φ85 lysate of aTB243 was used to transduce *ΔfacZ::specR* into aTB492 |
| **aTB515** | HG003 | *pTP63-gpsB-mNeon* | This study | Φ85 lysate of AGS063 was used to transduce *pTP63-gpsB-mNeon* into aTB003 |
| **aTB517** | HG003 | *pTP63-gpsB-mNeon pLOW-01855-mCherry* | This study | Φ85 lysate of aTB316 was used to transduce *pLOW-facZ-mCherry* into aTB515 |
| **aTB519** | HG003 | *pTP63-gpsB-mNeon ΔfacZ::specR* | This study | Φ85 lysate of aTB243 was used to transduce *ΔfacZ::specR* into aTB515 |
| **aTB521** | HG003 | *pKK30_RFP* | This study | Φ85 lysate of RN4220 *pKK30_RFP* was used to transduce pKK30_RFP into aTB003 |
| **aTB525** | HGoo3 | *ΔgpsB::Kan* | This study | Φ85 lysate of TAS201 was used to transduce *ΔgpsB::Kan* into aTB003 |
| **aTB527** | HG003 | *ΔfacZ::specR pKK30_RFP* | This study | Φ85 lysate of aTB521 was used to transduce pKK30_RFP into aTB251 |
| **aTB529** | HG003 | *ΔgpsB::kanR pKK30_RFP* | This study | Φ85 lysate of aTB521 was used to transduce pKK30_RFP into TAS201 |
| **aTB540** | HG003 | *ΔfacZ::specR pTP63-facZ ΔgpsB::kanR* | This study | Φ85 lysate of TAS201 was used to transduce *ΔgpsB::kanR* into aTB372 |
| **aTB542** | HG003 | *ΔfacZ::specR ΔgpsB::kanR pKK30-RFP* | This study | Φ85 lysate of aTB243 was used to transduce *ΔfacZ::specR* into aTB529 |
| **aTB549** | HG003 | *ΔfacZ::specR pTP63-facZ pKK30-RFP* | This study | Φ85 lysate of aTB521 was used to transduce *pKK30_RFP* into aTB372 |
| **aTP071** | RN4220 | pTP10 integrant | This study | pTP10 transformed and integrated into RN4220; cloning intermediate for Δ*atl* |
| **aTP103** | HG003 | Δ*atl* | This study | Φ85 lysate of aTP071 was used to transduce Δ*atl::kanR* into HG003 |
| **aTP436** | RN4220 | pTP044 pTP63-ezrA | This study | pTP63-ezrA transformed into RN4220 pTP044 |
| **aTP455** | RN4220 | pTP89 integrant | This study | pTP10 transformed and integrated into RN4220; cloning intermediate for Δ*ezrA* |
| **aTP481** | HG003 | *ΔezrA pTP63-ezrA* | This study | Unpublished (Rudner collection) |
| **TAS201** | RN4220 | *ΔgpsB::Kan* | This study | Provided by Walker lab |
| **TAS079** | RN4220 | *pLOW-gpsB-FLAG* | This study | Provided by Walker lab |
| **AGS063** | RN4220 | *pTP63-gpsB-mNeon* | This study | Provided by Walker lab |
| **SB100** | RN4220 | *pKK30-RFP* | This study | Provided by Walker lab |

**Table S8: *B. subtilis* strains used in this study**

| ***B. subtilis* strain** | **Relevant Genotype** | **Source** | **Construction Notes** |
|---|---|---|---|
| bDR11 | *WT* | Youngman, P. et al.[5] | - |
| bDR2229 | *amyE::*$P_{spac}$*-ftsZ-gfp* | Ben-Yehuda, S. & Losick, R.[6] | - |
| bDR2637 | *sacA::Pveg-mCherry(phleo)* | Roney, I. & Rudner, D.[7] | - |
| bDR2660 | *sacA::Pveg-BFP(phleo)* | Roney, I. & Rudner, D.[7] | - |
| bDR2789 | *sacA::Pveg-GFP(phleo)* | Roney, I. & Rudner, D.[7] | - |
| bTB013 | Δ*facZ::ermR* | This study | Marker crossed from deletion library into bDR11 |
| bTB018 | *amyE::Pspac-ftsZ-gfp* Δ*facZ::ermR* | This study | Marker crossed from deletion library into bDR2229 |
| bTB039 | *sacA::Pveg-GFP(phleo)* Δ*facZ::erm* | This study | Marker crossed from deletion library into bDR2789 |
| bTB040 | *sacA::Pveg-BFP(phleo)* Δ*facZ::erm* | This study | Marker crossed from deletion library into bDR2660 |

**Table S9: Plasmids used in this study**

| Plasmid | Relevant genotype & description | Marker | Source | Cut Sites/ ITA | Oligos |
|---|---|---|---|---|---|
| ***pTM378*** | ts origin, HMAR1 C9 transposase | Kan | Wang, H. et al.[4] | - | - |
| ***pTM381*** | ts origin, HMAR1 C9 transposase (truncated) | Kan | Wang, H. et al.[4] | - | - |
| ***pTP044*** | L54a integrase-bearing plasmid | Tet | Pang, T. et al.[3] | - | - |
| ***pLOW-ftsZ-GFP*** | *ftsZ-gfp* under $P_{spac}$ promoter | Erm | Liew, A. et al.[8] | - | - |
| ***pGL485*** | High-copy plasmid constitutively expressing LacI | Cm | Liew, A. et al.[8] | - | - |
| ***pKK30*** | dsRed, constitutively expressed red fluorescence | Tmp | Rodriguez, M. et al. [9] | - | - |
| ***pLOW-facZ*** | *facZ* under $P_{spac}$ promoter | Erm | Synthesized | - | - |
| ***pLOW-facZ-mCherry*** | *facZ-mCherry* under $P_{spac}$ promoter | Erm | Synthesized | - | - |
| ***pLOW-gpsB-GFP*** | *gpsB-mNeon* under $P_{spac}$ promoter | Erm | Walker lab | - | - |
| ***pTP63-lacZ*** | *lacZ* under $P_{tet}$ promoter | Cm | Pang, T. et al.[3] | - | - |
| ***pTP63-ezrA*** | *ezrA* under $P_{tet}$ promoter | Cm | Rudner lab, unpublished | - | - |
| ***pTP63-facZ*** | *facZ* under $P_{tet}$ promoter | Cm | This study | KpnI/EcoRI | oTB561, 562 |
| ***pTP63-facZ-mCherry*** | *facZ-mCherry* under $P_{tet}$ promoter | Cm | This study | KpnI/EcoRI | oTB563, 564 |
| ***pTP63-gpsB-mNeon*** | *gpsB-mNeon-FLAG* under $P_{tet}$ promoter | Cm | This study | ITA | AGS-059 through -064 |
| ***pTP63-ftsZ-GFP*** | *ftsZ-GFP* under $P_{tet}$ promoter | Cm | This study | KpnI/EcoRI | oTB566, 568 |
| ***pMR91-ΔfacZ::specR*** | 1.5 kb flanking sequence of *facZ* interrupted with *specR* marker, constitutive mScarlett, ts origin | Erm, Spec | This study | ITA | oTB490, 491, 492, 493 |
| ***pLOW-gpsB-FLAG*** | *gpsB-FLAG* under $P_{spac}$ promoter | Erm | This study | SalI/BamHI | oTS079, 80 |
| ***pSUMO-FacZ 3x(127-146)*** | Purification plasmid for His-SUMO-tagged FacZ, residues 127-146 | Kan | This study | ITA | oRWB137, oRWB138 |
| ***pSUMO-GpsB (1-75)*** | Purification plasmid for His-SUMO-tagged GpsB, residues 1-75 | Kan | This study | ITA | oRWB123, oRWB124 |

All plasmids produced in the course of this study were made by isothermal assembly (ITA) or restriction digest and ligation using pairs of restriction enzymes, as noted in this table. Plasmids were isolated from and are stored in either DH5α or BL21(DE3) *E. coli* strains, and are available upon request.

**Table S10: Oligos used in this study**

| Oligo | Name | Sequence | Purpose |
|---|---|---|---|
| **oTB045** | SpecR Seq F3 | TGATTCCACGGTACCATTTCTTGC | Outward-facing sequencing primer for downstream homology arm of Δ*facZ::specR* |
| **oTB065** | SpecR Seq R4 | AGTGCTCCCTGatGTCgacc | Outward-facing sequencing primer for upstream homology arm of Δ*facZ::specR* |
| **oTB169** | pMR091 MCS seq F | GACTTTACGAAACACGGAAACCG | Inward-facing sequencing primer for upstream homology arm of Δ*facZ::specR* |
| **oTB170** | pMR091 MCS seq R | atcagttcattgctcacgatatgtg | Inward-facing sequencing primer for downstream homology arm of Δ*facZ::specR* |
| **oTB192** | EzrA cPCR F | CATCTTCAATAAGGCTTGCTGC | Colony PCR for *ezrA* deletion |
| **oTB193** | EzrA cPCR R | CATCAGTCCAATTTGACAGAGTGC | Colony PCR for *ezrA* deletion |
| **oTB410** | pTP63_cPCR_F | CTCATTAAGCAGCTCTAATGCGC | Colony PCR for *pTP63* insert |
| **oTB411** | pTP63_cPCR_R | CCAGCGTTTCTGGGTGAGC | Colony PCR for *pTP63* insert |
| **oTB416** | del_Atl_cPCR_F1 | GGCGAAGTCGGCAAATACTTCG | Colony PCR for *atl* deletion |
| **oTB417** | del_Atl_cPCR_R1 | CGACGCATATCGTTGTAACACG | Colony PCR for *atl* deletion |
| **oTB490** | pMR091 01855 UpF | CGCTCGCGTATCGGTGATGGATCCCGATCAATTAGAGCAACTCGGTTATGTTTCG | Generate upstream flanking region of *facZ* for ITA |
| **oTB491** | pMR091 01855 UpR | cccggaaaaagagttgactaaatcaaTAAAAACGCCTCCTAATTAACATGTAATAATGTC | Generate upstream flanking region of *facZ* for ITA |
| **oTB492** | pMR091 01855 DnF | ccggtctatgttcatttagtcctccactaTTAATAATTAAACAAATGCACTTAAATGAGGTTGTTAC | Generate downstream flanking region of *facZ* for ITA |
| **oTB493** | pMR091 01855 DnR | gctctatataaaatatactcaaaatattatCCATGGtatGAATTCGCATTTGTTGTTTTTGAATTCTTATTACTA GTTTG | Generate downstream flanking region of *facZ* for ITA |
| **oTB494** | 01855 up out F | GATGATGTTTGTGCATTTATGGTTAATGAAGG | Δ*facZ* sequencing primer |
| **oTB495** | 01855 up in F | GATAATGCTGTTATTTTATTTATGGGTGCAGG | Δ*facZ* sequencing primer |
| **oTB496** | 01855 dn in R | GTTGCTGCGTCATTTGTATATCCTCC | Δ*facZ* sequencing primer |
| **oTB497** | 01855 dn out R | CATAGCTTACTGCTATGATTGATTATTCAACG | Δ*facZ* sequencing primer |
| **oTB498** | Spec Out Seq R | CAATAAACCCTTGCATAGGGATAACTTCG | Δ*facZ* sequencing primer |
| **oTB499** | Spec Out Seq F | CGTTACGTTATTAGTTATAGTTATTATAACATGTATTCACG | Δ*facZ* sequencing primer |
| **oTB500** | Spec Out Seq R2 | GCAGTTCGTAGTTATCTTGGAGAGAATATTGAATG | Δ*facZ* sequencing primer |
| **oTB501** | Spec Out Seq F2 | GTGTAAACCTATTCATTGTTTTAAAAATATCTCTTGCC | Δ*facZ* sequencing primer |
| **oTB513** | 01855 outside f | GTTGATTTTATCAAACAACAAAGAGAACCGG | Colony PCR for *facZ* deletion |
| **oTB514** | 01855 outside r | GGTAAGCACCTGAATGCCTACC | Colony PCR for *facZ* deletion |
| **oTB538** | pLOW MCS seq f1 | GACTTTATCTACAAGGTGTGGC | Colony PCR *for pLOW* insert |

| | | | |
|---|---|---|---|
| **oTB539** | pLOW MCS seq f2 | atcctctagagtcaattgtgagcgc | Colony PCR for *pLOW* insert |
| **oTB535** | PolyG-1st-1 primer | GTGACTGGAGTTCAGACGTGTGCTCTTCCGATCTGGGGGGGGGGGGGGG | Tn-seq primer (first PCR) |
| **oTB536** | Mariner PCR1 | GCCATCTATGTGTCTAGAGAC | Tn-seq primer (first PCR) |
| **oTB537** | Rnd2_Staph_IL | AATGATACGGCGACCACCGAGATCTACACTCTTTCGGGGACTTATCAGCCAACCTG | Tn-seq primer (second PCR) |
| **oTB538** | pLOW MCS seq f1 | GACTTTATCTACAAGGTGTGGC | Colony PCR/sequencing primer for *pLOW* constructs |
| **oTB540** | pLOW MCS seq r1 | TTCAGGCTGCGCAACTGTTG | Colony PCR/sequencing primer for *pLOW* constructs |
| **oTB545** | pGL485 fwd | taatgtATCGATaataatggtttcttagacg | Colony PCR for *pGL485* |
| **oTB546** | pGL485 rev | tattatGTCGACagtcggcattatctc | Colony PCR for *pGL485* |
| **oTB561** | pTP063d-01855 f | acaTAAGGAGGaGGTACCatgGATTGGATTTTACCAATTGCTGG | Subclone *facZ* into pTP63 vector; KpnI cut site |
| **oTB562** | pTP063d-01855 r | CCACCTGGAATTCttaTTTATCTACTCTAGAAGTATAGCTATGATTTGCATCAGTTGC | Subclone *facZ* into pTP63 vector; EcoRI cut site |
| **oTB563** | pTP063d-01855-mCh f | AGGAGGaGGTACCatgGATTGGATTTTACCAATTGCTGG | Subclone *facZ-mCherry* into pTP63 vector; KpnI cut site |
| **oTB564** | pTP063d-01855-mCh r | CACCTGGAATTCCttaGGATCCGCCAGCACCTTTG | Subclone *facZ-mCherry* into pTP63 vector; EcoRI cut site |
| **oTB566** | FtsZ-GFP KpnI F | GGAGGaGGTACCATGTTAGAATTTGAACAAGGATTTAATCATTTAGCG | Subclone *ftsZ-GFP* into pTP63 vector; KpnI cut site |
| **oTB568** | FtsZ-GFP EcoRI R | GAACGTCTTTCTTCTCTATTTCTAATGAAGC | Subclone *ftsZ-GFP* into pTP63 vector; EcoRI cut site |
| **oTB595** | GpsB cPCR F | GTTCTTCAAGCAGATGTTAGTTGATTTTATGG | Colony PCR for inactivation of *gpsB* |
| **oTB596** | GpsB cPCR R | GGACTTTCCTCTATATAATATAGCGATTACCC | Colony PCR for inactivation of *gpsB* |
| **oTB597** | GpsB Seq F | GTGATAAATTAAAAAATGTAGGAGGCGTCC | Colony PCR for inactivation of *gpsB* |
| **oTB598** | GpsB Seq R | CTTTCTTAGTTATCGCCTGACAATCTGGC | Colony PCR for inactivation of *gpsB* |
| **oTB599** | GpsB cPCR F | GGATAAAACAAACTATACTTGTGATATTGTG | Colony PCR for inactivation of *gpsB* |
| **oTB600** | GpsB cPCR R | CAATCATCTCAGACTGTGTGAGC | Colony PCR for inactivation of *gpsB* |
| **AGS-59** | pTP63 backbone F | GAATTCCAGGTGGCACTTTTCG | Amplification of pTP63 |
| **AGS-60** | pTP63 backbone R | GGTACCATCATACTCTATCAATGA | Amplification of pTP63 |
| **AGS-61** | GpsB F | GAGTATGATGGTACCgtttctaatgaggtggaaaaaATGTCAGATGTTTCATTGAAAT | Generate GpsB fragment for ITA of GpsB-mNeon |
| **AGS-62** | GpsB R | AGCTCCACCAGCGCTACCACCACCTTTACCAAATACAGCTTTTTCT | Generate GpsB fragment for ITA of GpsB-mNeon |

| | | | |
|---|---|---|---|
| **AGS-63** | mNeonGreen F | AGCGCTGGTGGAGCTGTGAGTAAAGGTGAGGAGGA | Generate mNeon fragment for ITA of GpsB-mNeon |
| **AGS-64** | mNeonGreen R | TGCCACCTGGAATTCTTActtgtcgtcatcgtctttgtagtcTTTATACAACTCATCCATTCCCAT | Generate mNeon fragment for ITA of GpsB-mNeon |
| **oTS079** | $rbs_{rpoB}$-GpsB F | GCAGGTCGACCATAATTTTTGAGGGGTGAATCTGTATGTCAGATGTTTCATTGAAATTATCA | Generate GpsB-FLAG insert |
| **oTS080** | GpsB-FLAG | CCCGGGGATCCTTACTTGTCGTCATCGTCTTTGTAG | Generate GpsB-FLAG insert |
| **AMo67** | gpsB_Wanner_KO_30H_U_o 67 | AAGATCAAAGTTTCTAATGAGGTGGAAAAA CATAAAAACAACTCGTAGCTTATCAAAG | Generate fragment for amplifying KanR marker (for deleting *gpsB*) |
| **AMo68** | gpsB_Wanner_KO_30H_D_o 68 | GTAAGACAGTTAAACTTTTGTATTTAGTAA CAATGACCTAAGAGGTGTGG | Generate fragment for amplifying KanR marker (for deleting *gpsB*) |
| **oRWB123** | GpsB SUMO Fwd | TGAACAGATTGGCGGCATGTCAG | Insert for pSUMO-FacZ 3x(127-146) |
| **oRWB124** | GpsB 75 SUMO Rev | catggatccttattacgtagcaacacgaaggcgaag | Insert for pSUMO-FacZ 3x(127-146) |
| **oRWB125** | Sumo-FacZ 127 Fwd | TgaacagattggcggcGAAATTGCAGATAAGTGGCAA | Insert for pSUMO-GpsB(1-75) |
| **oRWB126** | Sumo-FacZ 145 Rev | ccatggatccttattaTGCCTTGTAGTTTGCAGATCC | Insert for pSUMO-GpsB(1-75) |

### References


1 Omasits, U., Ahrens, C. H., Müller, S. & Wollscheid, B. Protter: interactive protein feature visualization and integration with experimental proteomic data. *Bioinformatics* **30**, 884-886, doi:10.1093/bioinformatics/btt607 (2014).

2 Fey, P. D. *et al.* A Genetic Resource for Rapid and Comprehensive Phenotype Screening of Nonessential Staphylococcus aureus Genes. *mBio* **4**, e00537-00512, doi:doi:10.1128/mBio.00537-12 (2013).

3 Pang, T., Wang, X., Lim, H. C., Bernhardt, T. G. & Rudner, D. Z. The nucleoid occlusion factor Noc controls DNA replication initiation in Staphylococcus aureus. *PLOS Genetics* **13**, e1006908, doi:10.1371/journal.pgen.1006908 (2017).

4 Wang, H., Claveau, D., Vaillancourt, J. P., Roemer, T. & Meredith, T. C. High-frequency transposition for determining antibacterial mode of action. *Nature Chemical Biology* **7**, 720-729, doi:10.1038/nchembio.643 (2011).

5 Youngman, P. J., Perkins, J. B. & Losick, R. Genetic transposition and insertional mutagenesis in Bacillus subtilis with Streptococcus faecalis transposon Tn917. *Proc Natl Acad Sci U S A* **80**, 2305-2309, doi:10.1073/pnas.80.8.2305 (1983).

6 Ben-Yehuda, S. & Losick, R. Asymmetric cell division in B. subtilis involves a spiral-like intermediate of the cytokinetic protein FtsZ. *Cell* **109**, 257-266, doi:10.1016/s0092-8674(02)00698-0 (2002).

7 Roney, I. J. & Rudner, D. Z. Two broadly conserved families of polyprenyl-phosphate transporters. *Nature* **613**, 729-734, doi:10.1038/s41586-022-05587-z (2023).

8 Liew, A. T. F. *et al.* A simple plasmid-based system that allows rapid generation of tightly controlled gene expression in Staphylococcus aureus. *Microbiology* **157**, 666-676, doi:https://doi.org/10.1099/mic.0.045146-0 (2011).

9 Rodriguez, M. D., Paul, Z., Wood, C. E., Rice, K. C. & Triplett, E. W. Construction of Stable Fluorescent Reporter Plasmids for Use in Staphylococcus aureus. *Frontiers in Microbiology* **8**, doi:10.3389/fmicb.2017.02491 (2017).
